# NECTIN4 defines a subtype-dependent tumor state coupling enfortumab vedotin targetability to antitumor immunity in bladder cancer

**DOI:** 10.64898/2026.10.07.757381

**Authors:** Kazuki Nishimura, Ryoichi Maenosono, Takuya Owari, Moritoshi Sakamoto, Takahiro Nishino, Kensuke Hirosuna, Takuya Tsujino, Takuya Izumi-Tamura, Yuki Yoshikawa, Miyu Azuma, Sakura Inoue, Asuka Kawachi, Atsuya Ishida, Kei Nishimori, Junko Mukohyama, Natsuko Shinohara, Marimu Sakumoto, Masataka Amisaki, Kazuki Okuyama, Mitsuaki Ishida, Masahiko Ajiro, Yoshinobu Hirose, Yutaka Suzuki, Hiroyoshi Nishikawa, Haruhito Azuma, Filipe L.F. Carvalho, Adam S. Kibel, Li Jia, Shohei Koyama, Takayuki Yoshino, Kazumasa Komura, Akihide Yoshimi

**Author notes:** Corresponding authors: Kazumasa Komura, MD, PhD, Professor, Department of Urology, Kawasaki Medical School, Address: 577 Matsushima, Kurashiki, Okayama 701-0192, Japan; Akihide Yoshimi, MD, PhD, Chief, Division of Cancer RNA Research, National Cancer Center Research Institute, Address: 5-1-1 Tsukiji, Chuo-ku, Tokyo 104-0045, Japan. These authors contributed equally (Kazuki Nishimura, Ryoichi Maenosono).

## Abstract

NECTIN4 is the target of enfortumab vedotin (EV), but its biological role in bladder cancer (BC) and its relationship to EV plus PD-1 blockade remain unclear. Integrating clinical multi-omics, spatial and single-cell immune profiling, and experimental models, we identify NECTIN4 as a subtype-dependent determinant of tumor state and antitumor immunity. In basal BC cells, NECTIN4 suppressed inflammatory and mesenchymal programs. Consistently, NECTIN4-high basal/squamous muscle-invasive BCs showed favorable survival, higher checkpoint inhibitor response rates, and a coordinated CD8+ T cell-inflamed microenvironment with no comparable associations in luminal papillary tumors. In orthotopic models, murine Nectin4 promoted CD8+ T cell-dependent tumor control, whereas human NECTIN4 conferred EV sensitivity. EV also attenuated NECTIN4-dependent TIGIT binding. Conversely, NECTIN4-low models were preferentially sensitive to HDAC inhibition. NECTIN4 therefore couples antibody-drug conjugate targetability to a subtype-specific tumor–immune state permissive for checkpoint blockade, whereas the NECTIN4-low state defines a distinct inflammatory-mesenchymal state with an alternative therapeutic vulnerability.

**eTOC blurb:** Nishimura et al. show that NECTIN4, the target of enfortumab vedotin, defines subtype-dependent tumor–immune states in bladder cancer. In basal/squamous tumors, NECTIN4 restrains inflammatory-mesenchymal programs and marks a CD8-rich microenvironment linked to checkpoint inhibitor response, whereas NECTIN4-low models show preferential sensitivity to HDAC inhibition.

**HIGHLIGHTS:**

- NECTIN4 is a subtype-dependent regulator of tumor state, not just an ADC target
- NECTIN4-high Ba/Sq tumors exhibit a coordinated CD8-inflamed microenvironment
- Nectin4 drives CD8 T cell-dependent tumor control in immunocompetent mice
- NECTIN4 couples EV targetability to a checkpoint-responsive immune context

## INTRODUCTION

Muscle-invasive bladder carcinoma (MIBC) is an aggressive malignancy with a high propensity for metastatic dissemination. Despite advances in systemic therapy, survival outcomes for patients with locally advanced or metastatic urothelial carcinoma (la/mUC) remain poor^1^.

The therapeutic landscape of urothelial carcinoma has been rapidly transformed by enfortumab vedotin (EV), a NECTIN4-targeted antibody–drug conjugate (ADC), combined with pembrolizumab. Following the striking survival benefit of EV plus pembrolizumab (EV+P) over platinum-based chemotherapy in first-line metastatic disease^2^, recent phase 3 trials have demonstrated substantial improvements in event-free and overall survival in the perioperative treatment of MIBC, across both cisplatin-ineligible and cisplatin-eligible populations^3,4^. Thus, EV+P is emerging as a therapeutic backbone across multiple stages of bladder cancer (BC). Despite this remarkable clinical activity, the biological basis for the interaction between NECTIN4-directed therapy and antitumor immunity remains poorly understood. In particular, it remains unclear whether NECTIN4 merely provides a cell-surface target for ADC delivery or also defines tumor-intrinsic and immune states that influence therapeutic response.

Although NECTIN4 has been implicated in diverse biological and clinical contexts across multiple cancer types^5–8^, its biological functions and clinicopathologic significance in bladder cancer (BC) remain incompletely understood. In parallel, large-scale transcriptomic studies have established that MIBC comprises distinct molecular subtypes characterized by divergent oncogenic programs, tumor microenvironmental states, and clinical behavior^9–12^. These observations raise the possibility that molecular subtype provides a critical context for understanding NECTIN4 biology and therapeutic relevance.

Here, we integrated multi-omics and spatial immune profiling of human BC with complementary in vitro and immunocompetent orthotopic models to define the tumor-intrinsic and immunologic functions of NECTIN4. We further asked whether these NECTIN4-defined states explain differential responsiveness to immune checkpoint blockade and shape the activity of EV plus PD-1 blockade and whether the NECTIN4-low state harbors alternative therapeutic vulnerabilities.

## RESULTS

### NECTIN4 expression is associated with favorable clinical outcomes in basal/squamous (Ba/Sq) MIBC

The OMPU-NCC urothelial carcinoma (UC) cohort comprised 469 UC patients and 35 normal bladder tissues. After excluding non-BC, 459 BC cases were included in the analysis (Figures 1A and S1A). None of the patients received EV during the follow-up period, excluding direct effects of EV exposure from these analyses.

**Figure 1.**
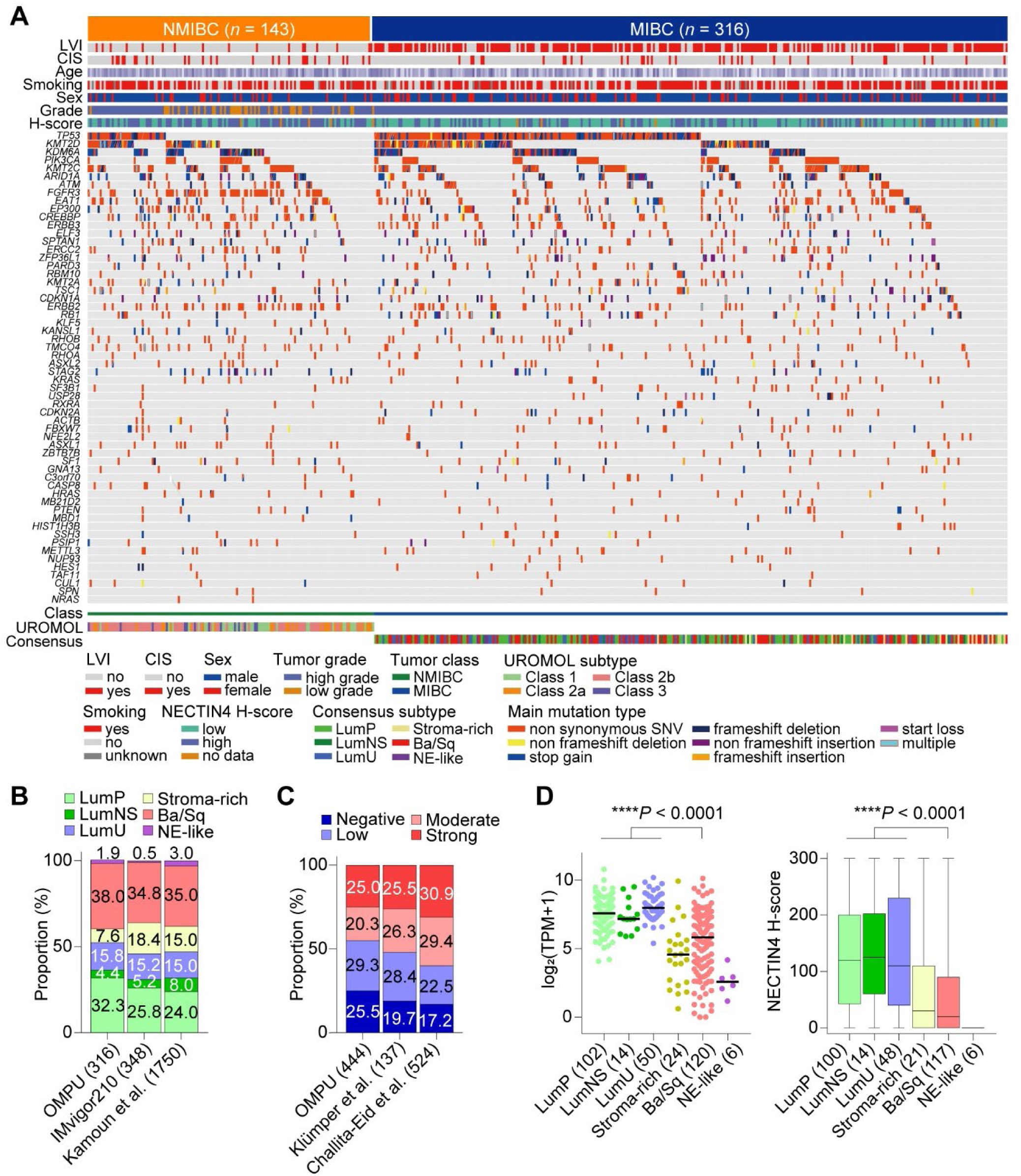
Clinicogenomic landscape and subtype-associated NECTIN4 expression in bladder cancer. (A) Clinicopathologic and genomic landscape of the OMPU-NCC cohort. (B) Consensus molecular subtype distributions in the indicated MIBC cohorts. (C) Distribution of NECTIN4 IHC categories: negative (H-score 0-14), low (15-99), moderate (100-199) and strong (200-300). (D) NECTIN4 mRNA expression and IHC H-scores across consensus molecular subtypes in the OMPU-NCC cohort. Horizontal lines indicate medians; boxplots show medians, interquartile ranges, and minimum-to-maximum whiskers. Numbers indicate tumors. *P* values were determined by Kruskal-Wallis test. \*\*\*\**P* < 0.0001. CIS, carcinoma in situ; LVI, lymphovascular invasion; MIBC, muscle-invasive bladder cancer; NMIBC, non-muscle-invasive bladder cancer; TPM, transcripts per million.

NECTIN4 protein expression was assessed by immunohistochemistry (IHC) on tissue microarrays (TMAs) in 444 patients, and H-scores positively correlated with NECTIN4 mRNA levels in both non-muscle-invasive bladder cancer (NMIBC) and MIBC (Figures S1B and S1C). Intratumoral heterogeneity of NECTIN4 staining was observed in a subset of cases (Figure S1B). Consensus molecular subtype for MIBC^12^ and the UROMOL 2021 molecular subtypes for NMIBC^13^ were assigned in 316 MIBC and 143 NMIBC cases from our cohort as well as the IMvigor210 cohort^14^. Consensus molecular subtype distributions in our cohort were broadly comparable to those in previously reported cohorts^12,14^, with modest cohort-specific differences (Figures 1B and S1D). Approximately one-fourth of tumors were NECTIN4-negative (H-score, 0-14), with a higher frequency in MIBC than in NMIBC (Figures 1C and S1E). Consistent with prior studies^15–17^, luminal MIBCs exhibited higher NECTIN4 levels than basal/squamous (Ba/Sq) MIBCs (Figure 1D and Figure S1F). Variant histology, such as sarcomatoid and small-cell carcinomas, showed particularly low NECTIN4 expression (Figure S1G).

Whole-exome sequencing with copy number analysis identified *NECTIN4* amplification in 14.5% of NMIBC cases and 16.3% of MIBC cases (Figure S1H). MIBCs with *NECTIN4* amplification exhibited significantly higher NECTIN4 protein expression (Figure S1I). Moreover, *NECTIN4* amplification was significantly enriched in luminal unstable (LumU) tumors relative to other MIBC subtypes (Figure S1J). Thus, NECTIN4 expression varies with molecular subtype and copy-number status.

We next examined clinicopathologic associations. NECTIN4 expression was markedly higher in BC than in normal bladder tissue, whereas MIBC showed significantly lower NECTIN4 expression than NMIBC (Figures 2A and 2B). Low NECTIN4 expression was associated with lymphovascular invasion, high-grade tumor, and smoking history (Figures S2A-S2E), while showing a modest positive correlation with age (Figure S2F). In intravesical recurrent tumors, NECTIN4 levels were largely comparable according to prior radiotherapy or chemotherapy exposure (Figures S2G and S2H).

**Figure 2.**
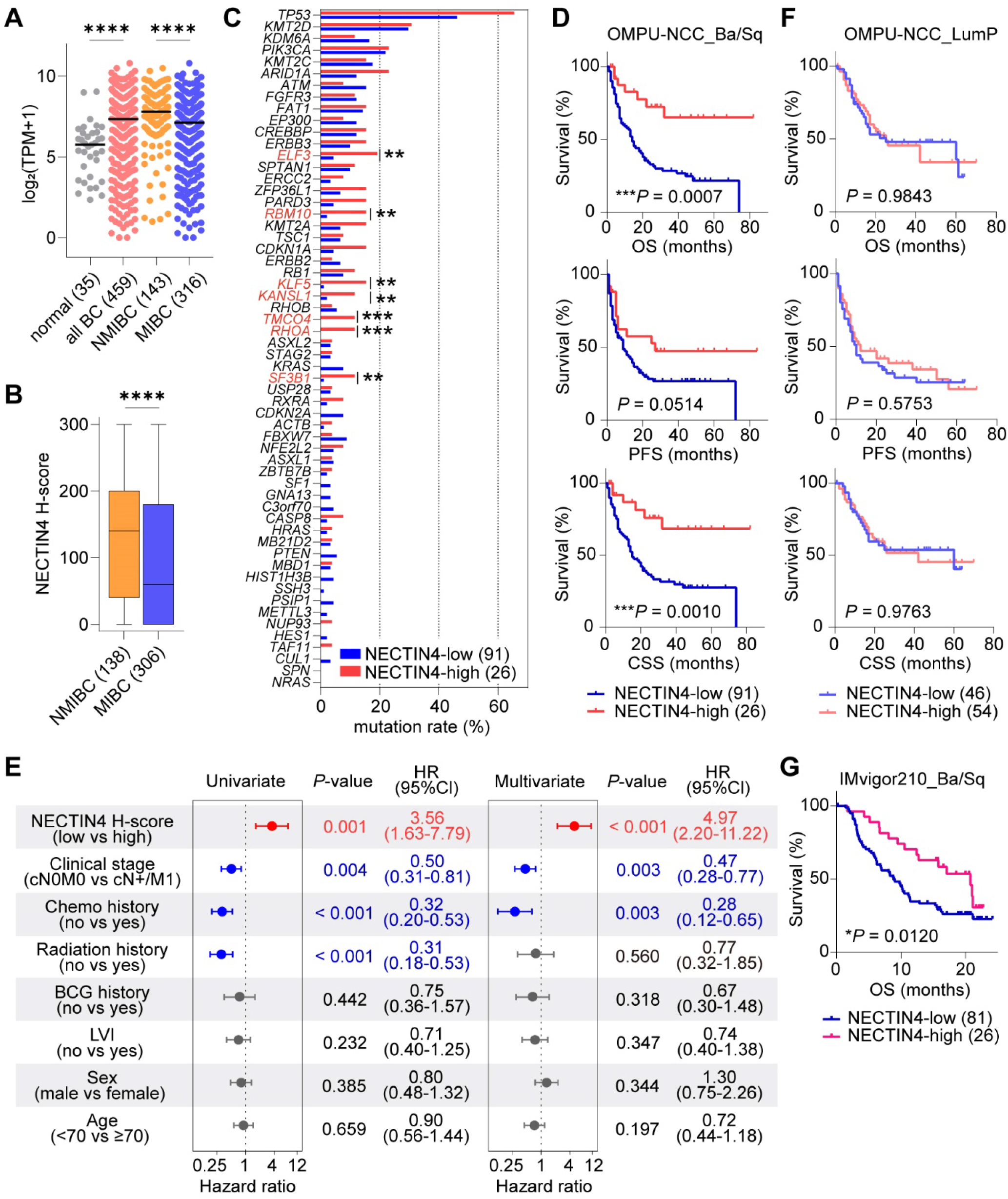
High NECTIN4 expression is associated with favorable outcomes in Ba/Sq MIBC. (A-F) Analyses of the OMPU-NCC cohort. (A) NECTIN4 mRNA expression in normal bladder tissues, all bladder cancers (BCs), non-muscle-invasive bladder cancers (NMIBCs), and muscle-invasive bladder cancers (MIBCs). (B) NECTIN4 IHC H-scores in NMIBC and MIBC. (C) Mutation frequencies in NECTIN4-low and -high Ba/Sq MIBCs. (D and F) Kaplan-Meier analyses of overall survival (OS), progression-free survival (PFS), and cancer-specific survival (CSS) in Ba/Sq (D) and LumP (F) MIBC. (E) Univariable and multivariable Cox proportional hazards analyses of OS in Ba/Sq MIBC. (G) Kaplan-Meier analysis of OS according to NECTIN4 expression in Ba/Sq tumors in the IMvigor210 cohort. Numbers indicate samples or patients. Horizontal lines indicate medians; boxplots show medians, interquartile ranges, and minimum-to-maximum whiskers. *P* values were determined by Mann-Whitney U test in (A) and (B), Fisher’s exact test in (C) and log-rank test in (D), (F), and (G). \**P* < 0.05, \*\**P* < 0.01, \*\*\**P* < 0.001 and \*\*\*\**P* < 0.0001. CI, confidence interval; HR, hazard ratio; LVI, lymphovascular invasion; TPM, transcripts per million.

We next asked whether NECTIN4 expression was associated with distinct biological and clinical features within individual molecular subtypes. For protein-level analyses, MIBCs were dichotomized at an H-score of 100, corresponding to the boundary between the low and moderate IHC categories defined above. We then focused on the two most prevalent subtypes, Ba/Sq and luminal papillary (LumP) (Figure S2I). Patient characteristics were largely comparable between NECTIN4-low and NECTIN4-high groups in both subtypes (Table S1).

Exploratory comparison of mutation frequencies identified several differences between NECTIN4-low and -high Ba/Sq tumors, whereas fewer differences were observed in LumP tumors (Figures 2C and S2J). Importantly, patients with NECTIN4-high Ba/Sq tumors had significantly longer overall survival (OS) and cancer-specific survival (CSS), with a trend toward longer progression-free survival (PFS) than those with NECTIN4-low tumors (Figure 2D). In multivariable Cox analysis, NECTIN4-high expression remained significantly associated with longer OS after adjustment for the clinical covariates included in the model (Figure 2E). In contrast, NECTIN4 expression was not prognostic in LumP tumors (Figure 2F). A similar subtype-specific survival association was observed in the independent IMvigor210 cohort, in which patients received anti-PD-L1 therapy: NECTIN4-high Ba/Sq tumors were associated with longer survival, whereas no significant association was observed in LumP tumors (Figures 2G and S2K). Moreover, an orthogonal genomic measure, NECTIN4 amplification, was also associated with longer survival in Ba/Sq but not LumP tumors (Figure S2L). Together, these findings support a subtype-specific association between high NECTIN4 expression and favorable clinical outcomes in Ba/Sq BC.

### NECTIN4 restrains growth, migration, and inflammatory programs in basal BC

Given the subtype-specific clinical association of NECTIN4, we investigated whether NECTIN4 directly influences tumor cell-intrinsic programs in basal BC. Consistent with prior reports^15^, basal BC cell lines expressed minimal NECTIN4, whereas luminal lines showed high expression (Figure S3A). To examine the consequences of restoring NECTIN4 in this low-expression context, we generated NECTIN4-overexpressing (OE) T24 and BOY-12E cell lines (Figures S3B and S3C). NECTIN4 overexpression significantly reduced colony formation and migration in both models (Figures 3A-3B and S3D), indicating that NECTIN4 can constrain growth- and motility-associated phenotypes in basal BC cells.

**Figure 3.**
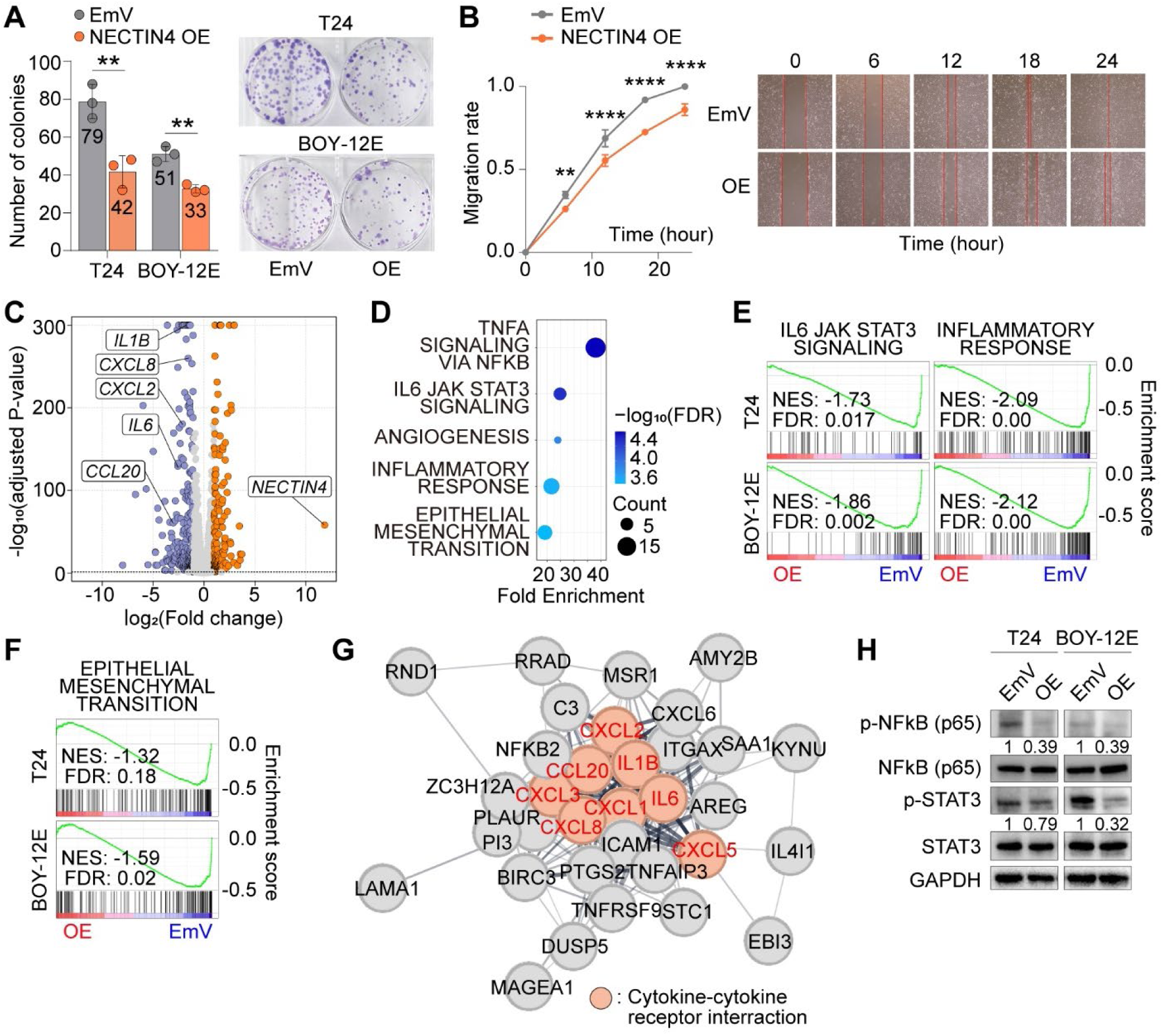
NECTIN4 suppresses growth, migration, and inflammatory-mesenchymal programs in basal bladder cancer cells. (A) Colony formation assays of empty vector (EmV) and NECTIN4-overexpressing (OE) cells. Numbers indicate median colony counts. (B) Scratch migration assay in T24 cells. (C) Differentially expressed genes between NECTIN4-OE and EmV T24 cells. (D) Top five Hallmark pathways enriched among genes commonly downregulated by NECTIN4 overexpression. (E-F) Gene set enrichment analysis of IL6-JAK-STAT3 signaling and inflammatory response (E) and epithelial-mesenchymal transition (F). (G) Protein-protein interaction network of commonly downregulated genes with a node degree of ≥3. Cytokine-cytokine receptor interaction genes are highlighted. (H) Immunoblot analysis of the indicated signaling proteins. Quantification represents phospho/total protein ratios normalized to the EmV control. Data are mean ± SD; *n* = 3 in (A) and (B). *P* values were determined by two-sided unpaired t test in (A) and two-way ANOVA in (B). \*\**P* < 0.01 and \*\*\*\**P* < 0.0001. FDR, false discovery rate; NES, normalized enrichment score.

Transcriptomic analysis identified 465 and 1440 differentially expressed genes between NECTIN4-OE and empty-vector (EmV) T24 and BOY-12E cells, respectively (Figure 3C and S3E-S3F). Among these, 56 genes were commonly downregulated and 24 genes were commonly upregulated across both models upon NECTIN4 overexpression (Figure S3G). Hallmark enrichment analysis of commonly downregulated genes highlighted inflammatory and mesenchymal programs, including TNFα/NFκB and IL6-JAK-STAT3 signaling, as well as epithelial-mesenchymal transition (EMT) (Figure 3D), whereas commonly upregulated genes were enriched for estrogen-response signatures (Figure S3H). Gene set enrichment analysis further showed reduced enrichment of inflammatory and EMT-associated signatures in NECTIN4-OE cells (Figures 3E and 3F). Protein-protein interaction analysis further identified IL6 and multiple CXCL/CCL family members as central nodes within the network of genes downregulated by NECTIN4 overexpression (Figure 3G). Consistent with these transcriptomic findings, immunoblotting demonstrated reduced NFκB and STAT3 phosphorylation in NECTIN4-OE cells (Figure 3H). These findings indicate that NECTIN4 suppresses NFκB/IL6-STAT3-associated inflammatory programs and restrains EMT-associated transcriptional states in basal BC cells.

To assess whether the transcriptional consequences of NECTIN4 perturbation differed in a luminal context, we established NECTIN4 knockdown (KD) in the luminal cell line KMBC-2 (Figures S3I and S3J). Transcriptomic profiling revealed enrichment of interferon-response and inflammatory-response programs following NECTIN4-KD (Figures S3K and S3L), whereas cell growth was minimally affected (Figure S3M). Thus, NECTIN4 perturbation had context-dependent effects, with the basal growth, NFκB/IL6-STAT3, and EMT phenotypes not recapitulated to the same extent in the luminal model.

### NECTIN4-low Ba/Sq tumors exhibit IL6/NFκB-associated inflammatory and mesenchymal programs

To determine whether the NECTIN4-associated transcriptional programs identified in basal cell models were recapitulated in clinical tumors, we analyzed RNA-seq profiles from 459 BC specimens in our cohort. In Ba/Sq MIBC, differential expression analysis (DEA) between NECTIN4-low and -high tumors revealed marked upregulation of inflammatory mediators in the NECTIN4-low group, with IL6 and CXCL8 among the most prominently upregulated genes (Figure 4A).

**Figure 4.**
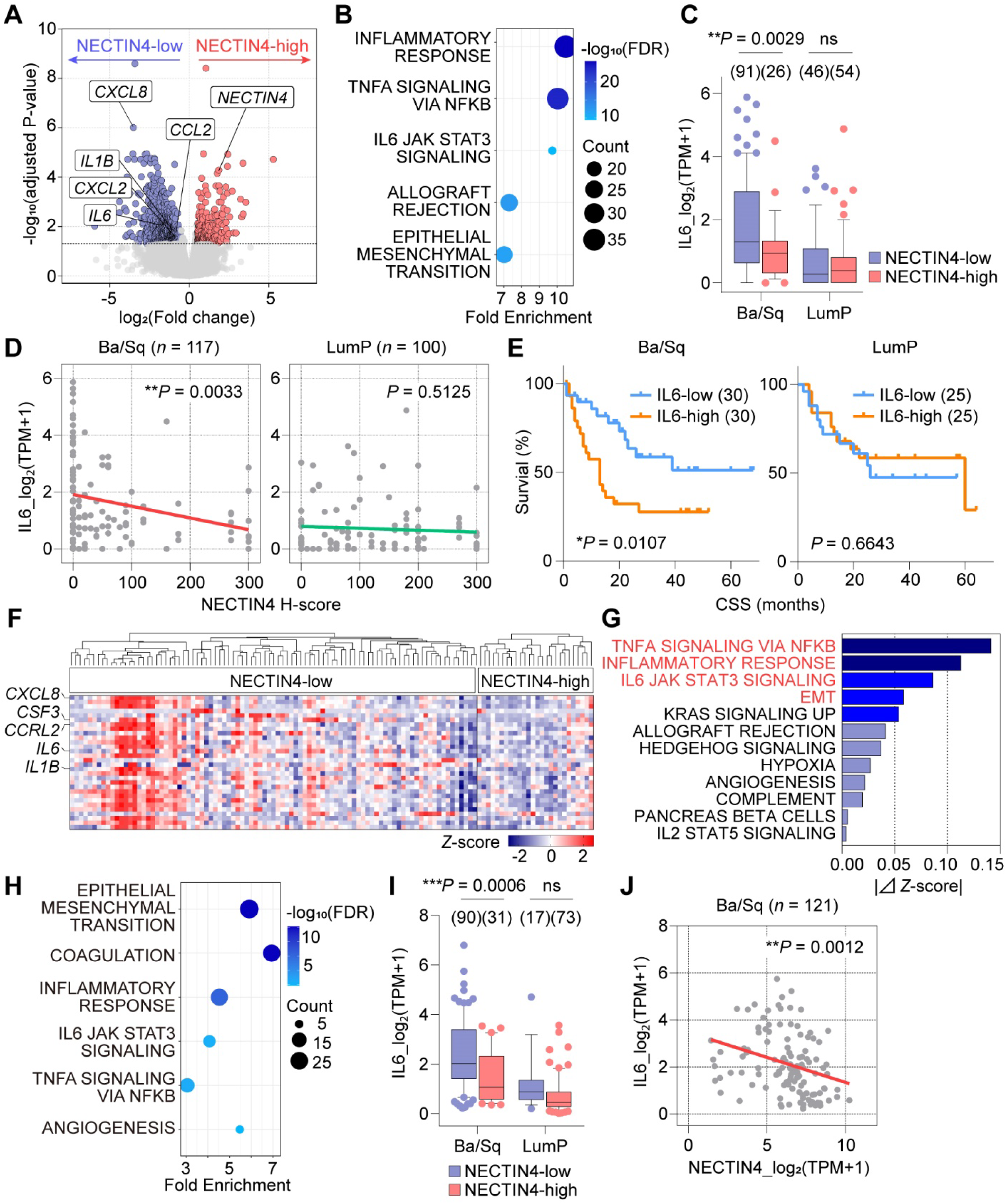
NECTIN4-low Ba/Sq tumors exhibit IL6/NFκB-associated inflammatory and epithelial-mesenchymal transition (EMT) programs. (A-G) Analyses of the OMPU-NCC cohort. (A) Differentially expressed genes between NECTIN4-low and -high Ba/Sq MIBCs. (B) Top five Hallmark pathways enriched in NECTIN4-low Ba/Sq tumors. (C) IL6 expression according to NECTIN4 status and subtype. (D) Correlations between NECTIN4 H-score and IL6 expression. (E) Kaplan-Meier analyses of cancer-specific survival (CSS) comparing tumors in the highest and lowest quartiles of IL6 expression within each subtype. (F) Heatmap showing the top 30 inflammation-related genes upregulated in NECTIN4-low Ba/Sq tumors. (G) Differences in mean Hallmark pathway z-scores between NECTIN4-low and -high Ba/Sq tumors. Pathways with higher scores in NECTIN4-low tumors are shown. (H-J) Analyses of the IMvigor210 cohort. (H) Enrichment of inflammation- and EMT-related Hallmark pathways in NECTIN4-low Ba/Sq tumors. (I) IL6 expression according to NECTIN4 status and subtype. (J) Correlation between NECTIN4 and IL6 expression in Ba/Sq tumors. Numbers indicate tumors. *P* values were determined by Mann-Whitney U test in (C) and (I), log-rank test in (E), and simple linear regression in (D) and (J). \**P* < 0.05, \*\**P* < 0.01 and \*\*\**P* < 0.001; ns, not significant.

Hallmark enrichment analysis showed robust enrichment of inflammatory and EMT-associated programs, including TNFα/NFκB and IL6-JAK-STAT3 signaling, in NECTIN4-low Ba/Sq tumors (Figure 4B). In contrast, LumP MIBCs exhibited substantially fewer transcriptomic differences between NECTIN4-low and -high groups and did not show comparable enrichment of these inflammatory and EMT-associated programs (Figure S4A). Consistently, IL6 and CXCL8 expression levels were significantly higher in NECTIN4-low Ba/Sq tumors but not in LumP tumors (Figure 4C and Figure S4B). NECTIN4 H-scores were inversely associated with IL6 mRNA levels specifically in Ba/Sq tumors, whereas no significant association was observed in LumP tumors (Figure 4D). Clinically, high IL6 expression was associated with shorter CSS in Ba/Sq but not LumP tumors (Figure 4E), consistent with the adverse clinical features of the IL6-high inflammatory state.

Heatmap visualization and sample-level Hallmark pathway scoring further showed that the largest transcriptional differences between NECTIN4-low and -high Ba/Sq tumors were dominated by inflammatory and EMT-associated programs (Figures 4F and 4G). Notably, the elevated inflammatory pathway scores were largely restricted to NECTIN4-low Ba/Sq tumors (Figure S4C). Consistent with the EMT-associated transcriptional signature, NECTIN4-low Ba/Sq tumors showed lower expression of the epithelial marker CDH1 and higher expression of the mesenchymal marker CDH2. Moreover, NECTIN4 H-score was positively associated with CDH1 and inversely associated with CDH2 expression, while these associations were not evident in LumP tumors (Figures S4D and S4E).

We next analyzed transcriptomic data from the IMvigor210 cohort to assess these associations in an independent dataset. NECTIN4-low Ba/Sq tumors similarly exhibited enrichment of TNFα/NF-κB-, IL6-JAK-STAT3-, inflammatory response-, and EMT-associated pathways (Figure 4H and S4F), elevated IL6 expression (Figure 4I), and inverse correlations between NECTIN4 and both IL6 or CDH2, together with a positive association with CDH1 (Figure 4J and S4G-S4H). By contrast, in LumP tumors, NECTIN4 status was not associated with significant differences in IL6, CDH1, or CDH2 expression, and only the Hallmark inflammatory response signature was enriched among the inflammation- and EMT-related programs examined (Figure 4I and S4G-S4I). Together, these findings identify NECTIN4-low Ba/Sq MIBC as an IL6/NFκB-high, inflammatory-mesenchymal transcriptional state associated with unfavorable clinical outcomes.

### NECTIN4-high Ba/Sq MIBC is associated with a coordinated T cell-inflamed tumor microenvironment

To distinguish the inflammatory-mesenchymal state enriched in NECTIN4-low tumors from productive antitumor immune inflammation, we investigated whether NECTIN4 expression was associated with immune-cell composition and clinical response to immune checkpoint inhibitors (ICIs)^18–22^. To minimize treatment-related confounding, analyses of the tumor microenvironment (TME) were restricted to treatment-naïve MIBC cases (Figure S5A). NECTIN4-high treatment-naïve Ba/Sq tumors remained associated with longer CSS than NECTIN4-low tumors (Figure S5B).

CIBERSORTx deconvolution in absolute mode estimated immune-cell abundance. NECTIN4-high tumors showed significantly greater inferred abundance of CD8+ T cells, resting memory CD4+ T cells, and regulatory T cells (Tregs) than NECTIN4-low tumors (Figure 5A). Immune-cell populations also exhibited distinct patterns of coordinated abundance according to NECTIN4 status, with broad positive co-abundance among T-cell populations in NECTIN4-high tumors (Figure 5B and Table S2). Higher estimated abundance of both CD8+ T cells and Tregs was significantly associated with longer CSS (Figure S5C). Their co-enrichment and positive covariation suggest that Treg abundance in this context accompanies a T cell-rich immune state rather than isolated immunosuppression. Consistently, CD8-high/Treg-high tumors were associated with longer CSS (Figure S5D). These findings support co-enrichment of effector and regulatory T-cell populations within a T cell-inflamed Ba/Sq microenvironment associated with favorable clinical outcomes. In contrast, LumP tumors showed no significant NECTIN4-associated differences in inferred immune-cell abundance, and the corresponding correlation matrices did not reveal a comparable NECTIN4-associated pattern (Figures S5E-S5F and Table S2).

**Figure 5.**
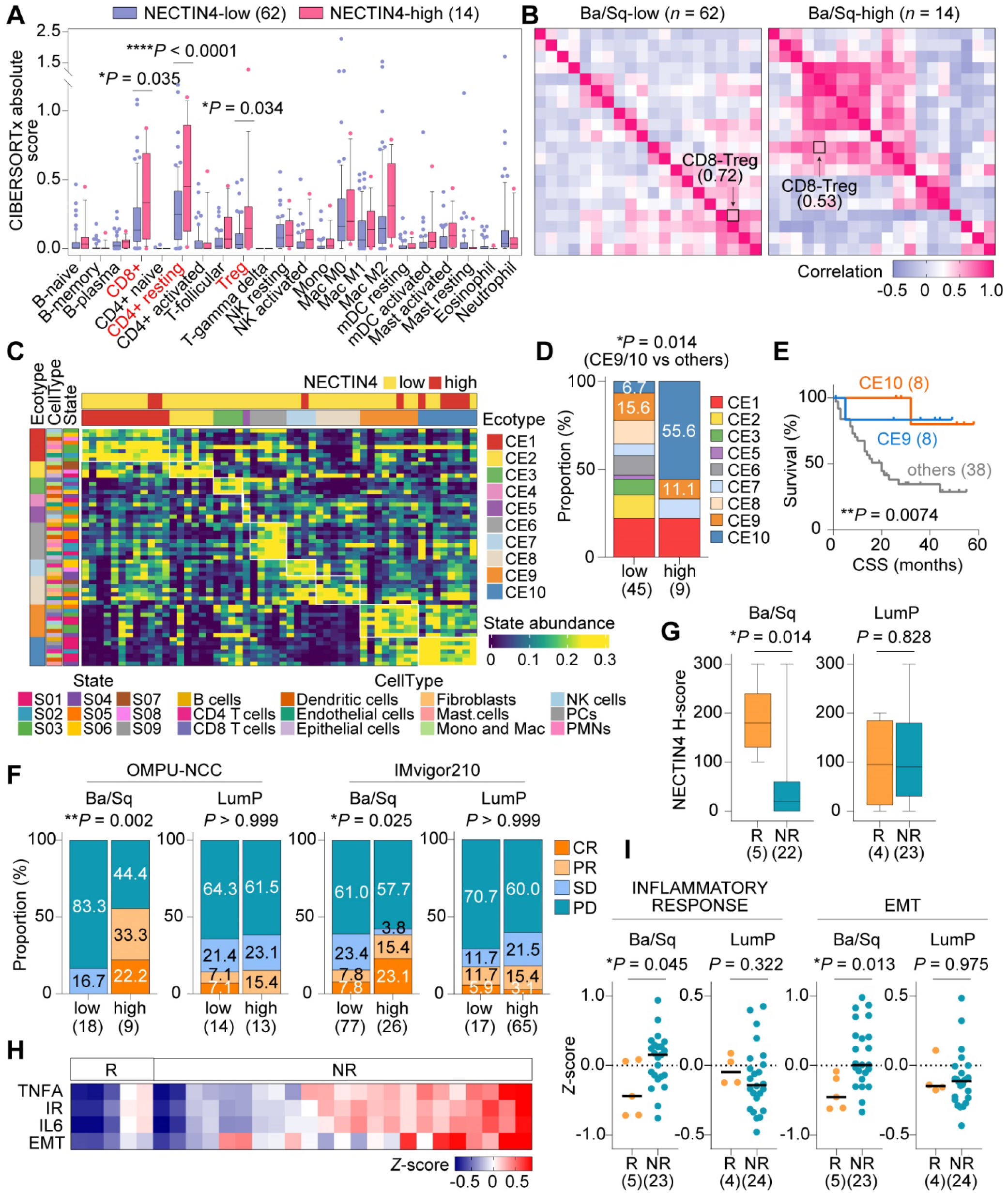
NECTIN4-high Ba/Sq tumors exhibit a coordinated T cell-inflamed TME and are associated with higher response rates to immune checkpoint inhibitors (ICIs) (A-E) Analyses of treatment-naïve Ba/Sq tumors in the OMPU-NCC cohort. (A) CIBERSORTx-inferred immune-cell abundance. (B) Pearson correlation matrices of inferred immune-cell abundance. (C-E) EcoTyper-derived cell states and cell ecotypes (CEs) (C), distribution by NECTIN4 status (D) and associated cancer specific survival (CSS) (E). (F) Clinical responses to ICIs according to NECTIN4 status and molecular subtype. (G-I) Analyses of ICI-treated patients in the OMPU-NCC cohort. (G) NECTIN4 H-scores in ICI responders and nonresponders. (H) Hallmark pathway z-scores in Ba/Sq responders and nonresponders. (I) Inflammatory response and EMT Hallmark pathway z-scores in responders and nonresponders. Responders (R) included complete response (CR) and partial response (PR); nonresponders (NR) included stable disease (SD) and progressive disease (PD). Numbers indicate patients. Horizontal lines indicate medians. Boxplots show medians and interquartile ranges, with 10th-90th percentile whiskers in (A) and minimum-to-maximum whiskers in (G). *P* values were determined by two-way ANOVA in (A), Fisher’s exact test in (D) and (F), log-rank test in (E), and Mann-Whitney U test in (G) and (I). \**P* < 0.05, \*\**P* < 0.01 and \*\*\*\**P* < 0.0001. Treg, Regulatory T cells; TNFA, TNFα signaling via NFκB; IR, inflammatory response; IL6, IL6-JAK-STAT3 signaling; EMT, epithelial-mesenchymal transition.

We next applied EcoTyper to characterize immune ecotypes^23^. NECTIN4-high Ba/Sq tumors were enriched for immune-inflamed CE9 and CE10 ecotypes, whereas CE6, which has been associated with poor immunotherapy response, was observed exclusively in NECTIN4-low tumors (Figures 5C-5D and S5G). Patients with CE9/CE10 Ba/Sq tumors had longer CSS than those with other ecotypes (Figure 5E). These associations were not observed in LumP tumors (Figures S5H and S5I).

Given these T cell-inflamed features, we evaluated ICI response according to NECTIN4 expression. In both our cohort and the IMvigor210 cohort, NECTIN4-high Ba/Sq tumors were associated with significantly higher objective response rates to anti-PD-1/PD-L1 therapy, whereas no association was observed in LumP tumors (Figure 5F). In the OMPU-NCC cohort, Ba/Sq responders had significantly higher NECTIN4 H-score than nonresponders (Figure 5G). In IMvigor210, NECTIN4 mRNA expression showed a similar trend but did not reach statistical significance (Figure S5J). Moreover, Ba/Sq responders showed lower inflammatory response and EMT signature scores than nonresponders, whereas no significant differences were observed in LumP tumors (Figures 5H and 5I). Together, these findings support a distinct NECTIN4-high tumor-immune state characterized by reduced inflammatory-mesenchymal programs, coordinated T-cell inflammation, and higher ICI response rates specifically within Ba/Sq MIBC.

### NECTIN4-high Ba/Sq MIBCs exhibit a spatially organized CD8-rich, TIGIT-expressing TME

To spatially resolve the immune architecture associated with NECTIN4 expression, we performed high-plex spatial immune profiling of Ba/Sq MIBC TMAs using CODEX-based multiplex IF (mIF) on the PhenoCycler-Fusion platform^24,25^ (Figure 6A). After quality control, 108 Ba/Sq cases (NECTIN4-low, *n* = 82; NECTIN4-high, *n* = 26) were analyzed. Unsupervised clustering identified four recurrent cellular regions with distinct TME features: an immune/extratumoral region (R1), a NECTIN4-negative tumor region (R2), a NECTIN4-positive tumor region (R3), and an αSMA-rich/vascular region (R4) (Figure 6B). As expected from the IHC classification, NECTIN4-high tumors contained a higher proportion of R3 than NECTIN4-low tumors (Figures 6C and S6A). Notably, the NECTIN4-positive R3 region was characterized by greater representation of CD8+ T cells, including PD-1+TIGIT+CD8+ T cells, as well as Tregs relative to the NECTIN4-negative R2 region (Figures 6B and 6D), indicating that NECTIN4-positive tumor regions are associated with a distinct local immune composition. Given the intratumoral heterogeneity of NECTIN4 staining in some tumors (Figure S1B), these findings further suggest that regional variation in NECTIN4 expression may accompany spatially distinct immune microenvironments within individual tumors.

**Figure 6.**
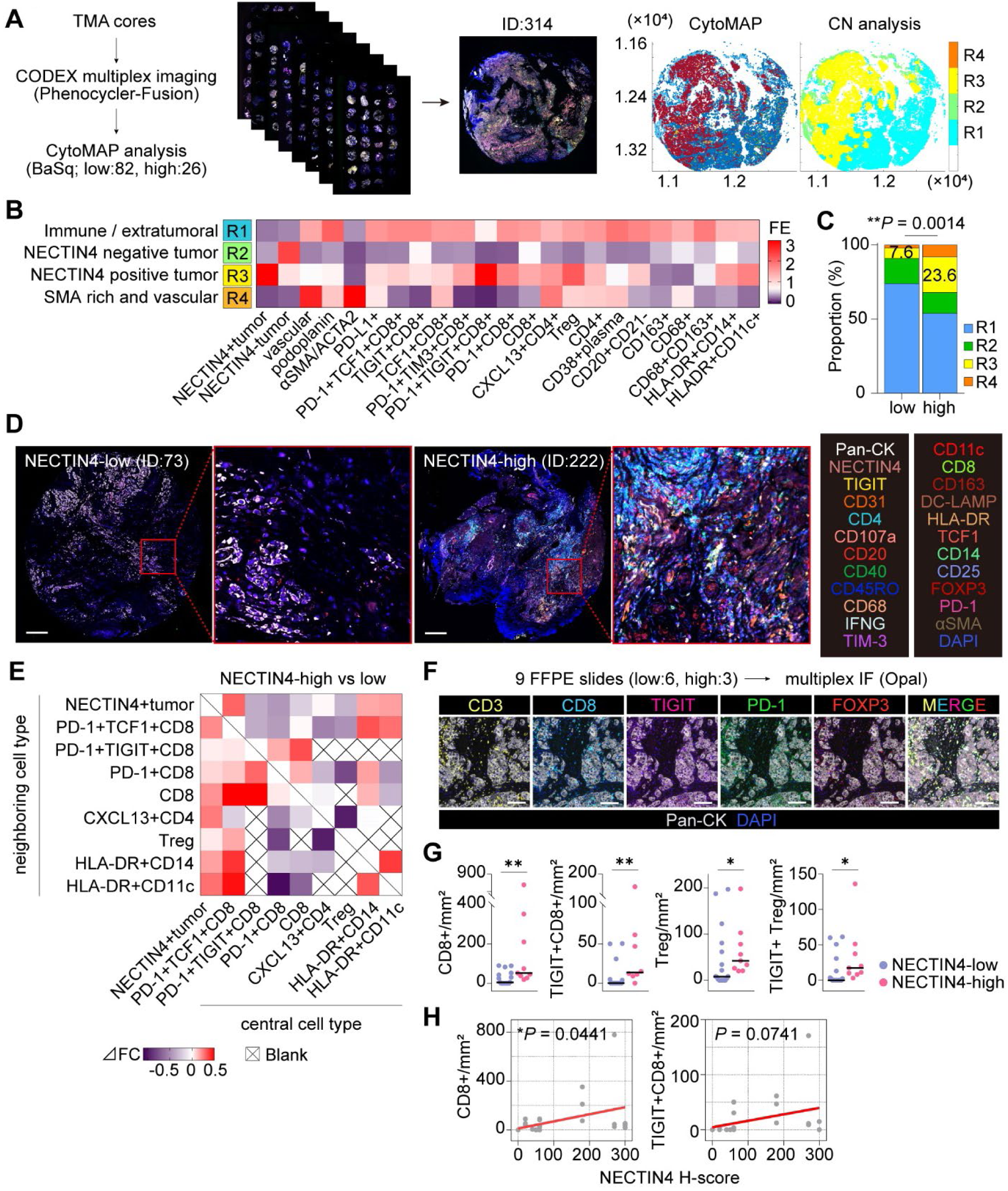
NECTIN4-high Ba/Sq MIBCs exhibit a CD8+ T-cell-enriched, TIGIT-expressing immune microenvironment. (A) Workflow for CODEX multiplex immunofluorescence (mIF) and CytoMAP-based analysis. (B) Fold enrichment (FE) of the indicated cellular phenotypes across the four identified regions. (C) Distribution of the four regions by NECTIN4 status. The proportion of R3 versus all other regions was compared between groups. (D) CODEX images of NECTIN4-low and -high tumors. Scale bar, 250 µm. (E) Differences in nearest-neighbor associations in NECTIN4-high versus -low tumors. Positive values indicate increased associations in NECTIN4-high tumors. (F) Opal-based mIF workflow and representative T-cell images. Scale bar, 100 µm. (G) Densities of CD8+ T cells, TIGIT+CD8+ T cells, regulatory T cells (Tregs) and TIGIT+ Tregs in NECTIN4-low (L) and -high (H) tumors. (H) Correlation between NECTIN4 H-score and CD8+ or TIGIT+CD8+ T-cell density. Horizontal lines indicate medians. *P* values were determined by Fisher’s exact test in (C), Mann-Whitney U test in (G), and simple linear regression in (H). \**P* < 0.05 and \*\**P* < 0.01. FFPE, formalin-fixed, paraffin-embedded; SMA, smooth muscle actin; TMA, tissue microarray.

Correlation analysis revealed broader positive associations among tumor and immune features in NECTIN4-high than in NECTIN4-low tumors (Figure S6B). Nearest-neighbor analysis further showed increased spatial associations between NECTIN4-positive tumor cells and several T-cell and myeloid-cell populations in NECTIN4-high tumors (Figure 6E).

To orthogonally assess these immune features using an independent imaging platform, we performed Opal-based mIF in a set of treatment-naïve Ba/Sq tumors using two complementary panels (Figures 6F and S6C). NECTIN4-high tumors showed higher densities of CD8+ T cells and Tregs across tumor and stromal compartments, with prominent enrichment of TIGIT+CD8+ T cells and TIGIT+ Tregs (Figures 6G and S6D-S6E). By contrast, PD-1+CD8+ T-cell density did not significantly differ according to NECTIN4 status in this validation set (Figures S6D and S6E). CD8+ T-cell density was significantly positively associated with NECTIN4 H-score, whereas TIGIT+CD8+ T-cell density showed a similar positive trend that did not reach statistical significance (Figure 6H). The intratumoral CD8/Treg ratio was also higher in NECTIN4-high tumors, whereas the stromal CD8/Treg ratio did not significantly differ between groups (Figure S6F). Major myeloid-cell populations were largely comparable between groups; however, stromal HLA-DR+CD11c+ dendritic cell (DC) density was higher in NECTIN4-high tumors (Figures S6D and S6E).

To further examine the cellular states associated with TIGIT expression in NECTIN4-high Ba/Sq tumors, we performed an integrative analysis of publicly available single-cell RNA-seq datasets from MIBC tumors^26,27^. Among the available datasets, five tumors were classified as Ba/Sq and stratified into NECTIN4-low (*n* = 2) and NECTIN4-high (*n* = 3) groups based on pseudo-bulk NECTIN4 expression. Following quality control and batch correction, CD4+ and CD8+ T cells were separately projected onto reference TIL atlases (Figures S6G and S6H). TIGIT-high CD4+ T cells were predominantly Tregs, whereas TIGIT-high CD8+ T cells comprised mainly effector-memory and exhausted states (Figures S6G and S6H). At the patient level, TIGIT-high cell fractions were descriptively higher in the three NECTIN4-high tumors than in the two NECTIN4-low tumors in both CD4+ and CD8+ compartments (Figure S6I). Together, these analyses associate high NECTIN4 expression with a spatially organized CD8-rich immune microenvironment enriched for TIGIT-expressing CD8+ T cells and Tregs in Ba/Sq MIBC.

### NECTIN4 promotes a CD8-rich TME and CD8+ T-cell-dependent tumor control

To determine whether NECTIN4 contributes to the CD8-rich TME observed in NECTIN4-high tumors, we established an immunocompetent orthotopic BC model using murine MB49 cells. Because parental MB49 cells expressed minimal endogenous Nectin4, we generated stable lines overexpressing either murine Nectin4 (mOE) or human NECTIN4 (hOE) (Figures S7A and S7B). We used mOE tumors to establish Nectin4-dependent immune phenotypes in a syngeneic setting, whereas hOE tumors provided an EV-targetable model for pharmacologic studies.

Flow-cytometric profiling two weeks after intravesical bladder implantation showed a trend toward increased total T-cell representation in mOE tumors (Figure S7C). Both CD8+ and CD4+ T-cell fractions were significantly increased in mOE compared with empty-vector (EmV) tumors (Figure 7A). The proportion of PD-1+TIGIT+CD8+ T cells was also elevated, consistent with enrichment of a checkpoint-expressing CD8+ T-cell population (Figure 7A). Treg frequencies showed a nonsignificant increase (Figure S7D), whereas major myeloid subsets were largely unchanged (Figure S7E). Parallel experiments using hOE tumors showed similar increases in total T cells, CD8+ T cells, and PD-1+TIGIT+CD8+ T cells (Figure S7F), indicating that the EV-targetable hOE model recapitulated the T-cell phenotype observed with murine Nectin4.

**Figure 7.**
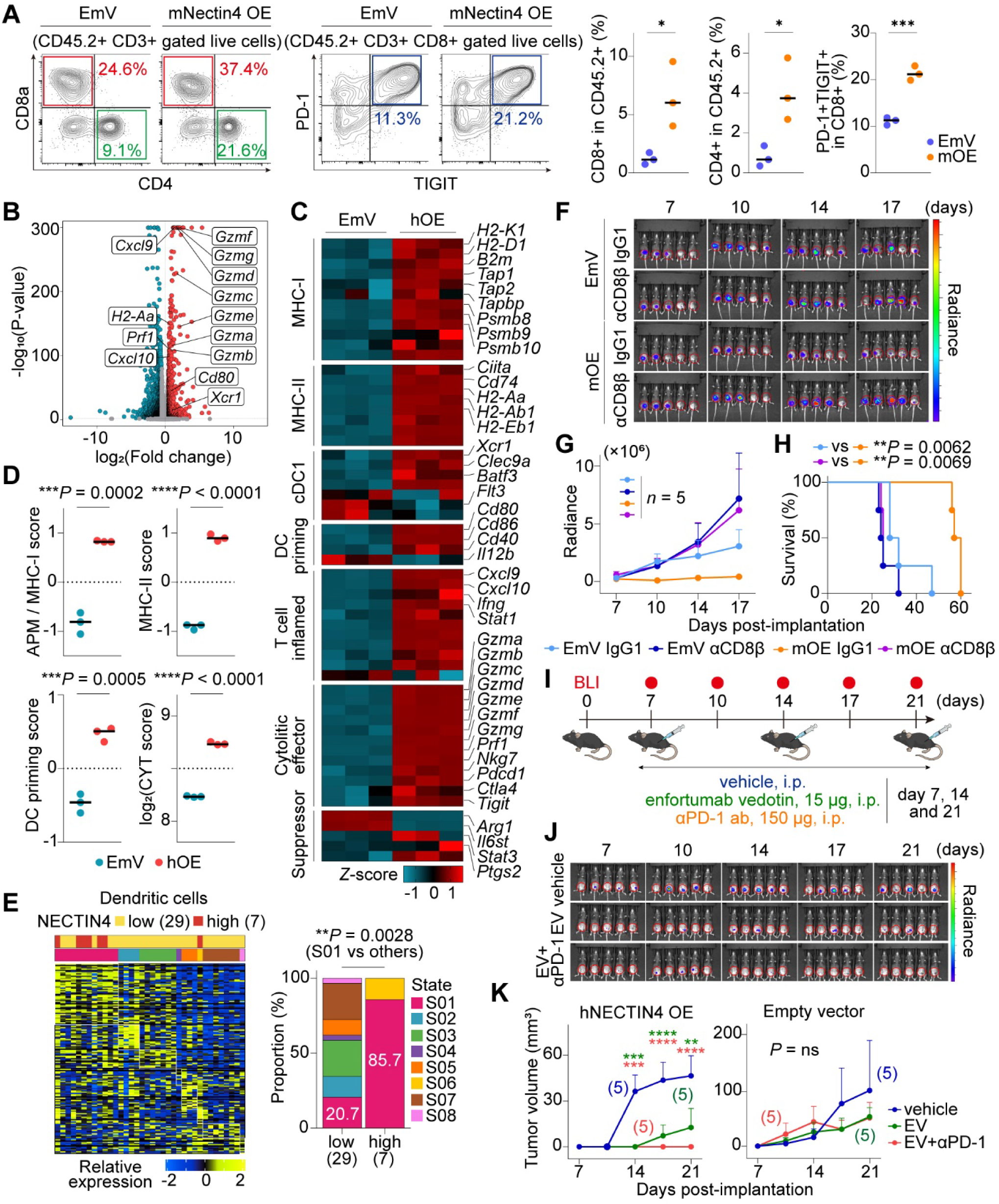
NECTIN4 promotes CD8+ T cell-dependent tumor control and supports the activity of EV plus PD-1 blockade. (A) Representative flow-cytometry plots of CD4+, CD8+, and PD-1+TIGIT+CD8+ T cells. Quantification shows CD4+ and CD8+ T-cell frequencies among CD45.2+ cells and PD-1+TIGIT+ frequencies among CD8+ T cells in orthotopic (EmV) and murine Nectin4-overexpressing (mOE) tumors. *n* = 3. (B-D) Differential gene expression (B), immune-program gene expression (C) and antigen-processing machinery/MHC-I (APM/MHC-I), MHC-II, DC-priming, and cytolytic activity (CYT) scores (D) in CD45.2+ cells from human NECTIN4-overexpressing (hOE) and EmV tumors. *n* = 3. (E) EcoTyper-derived DC-state distribution in NECTIN4-low and -high Ba/Sq MIBCs. (F-H) Bioluminescence images (BLI) (F), tumor radiance (G), and survival (H) of mice bearing orthotopic EmV or mOE tumors treated with IgG1 or anti-CD8β. *n* = 5 per group. (I-K) Treatment schema (I), BLI (J) and tumor growth (K) in orthotopic hOE and EmV tumors treated with vehicle, enfortumab vedotin (EV), or EV plus anti-PD-1 antibody (EV+αPD-1). Data in (G) and (K) are mean ± SEM. Numbers indicate patients in (E) and mice in (K). Horizontal lines indicate medians. *P* values were determined by two-sided unpaired t test in (A) and (D), Fisher’s exact test in (E), two-way ANOVA in (G) and (K), and log-rank test in (H). \**P* < 0.05, \*\**P* < 0.01, \*\*\**P* < 0.001 and \*\*\*\**P* < 0.0001; ns, not significant.

To characterize immune programs present in the EV-targetable hOE model, we performed transcriptomic analyses of CD45.2+ immune cells isolated from orthotopic hOE and EmV tumors (Figure 7B). Differential expression and pathway enrichment analyses revealed enrichment of antigen processing/presentation and immune-effector programs in immune cells from hOE tumors (Figures 7B and S7G). MHC class I components, antigen-processing machinery (APM), and MHC class II pathway genes were coordinately upregulated, resulting in higher composite APM/MHC-I and MHC-II scores (Figures 7B-7D). Genes associated with cross-presenting DC programs and costimulatory molecules were also upregulated, with higher DC-priming scores (Figures 7B-7D). Importantly, in human Ba/Sq MIBC, EcoTyper analysis identified enrichment of the priming-associated DC state S01 in NECTIN4-high tumors (Figure 7E). In addition, immune cells from hOE tumors exhibited increased expression of T-cell-inflamed chemokines, including Cxcl9 and Cxcl10, interferon-response genes, and cytotoxic effector molecules, including Prf1 and multiple Gzm family members, together with higher cytolytic activity scores (Figures 7B-7D). Tigit and Pdcd1 were also elevated, consistent with a checkpoint-expressing, T-cell-inflamed state. Together, these findings indicate that the hOE immune microenvironment is characterized by coordinated antigen presentation, DC-associated, T-cell-inflamed, and cytotoxic-effector programs that are also supported by immune features observed in human NECTIN4-high Ba/Sq tumors.

To determine whether CD8+ T cells were required for the tumor-control phenotype associated with mNectin4 expression, we performed CD8 depletion in orthotopic mOE and EmV tumors using an anti-CD8β antibody (Figure S7H). Efficient systemic CD8+ T-cell depletion was confirmed (Figure S7I). In isotype-treated mice, mOE tumors exhibited slower growth and significantly longer survival compared with EmV controls (Figures 7F-7H). However, CD8 depletion accelerated tumor progression and shortened survival in both models and substantially diminished the tumor-control advantage associated with mOE tumors (Figures 7F-7H). These results demonstrate that the tumor-control phenotype conferred by mNectin4 expression is dependent, at least in substantial part, on CD8+ T cells. Collectively, mOE tumors establish that mNectin4 can promote a CD8-rich TME and CD8+ T-cell-dependent tumor control whereas hOE transcriptomic data and human tumor data associate the NECTIN4-high state with antigen-presentation, DC-associated, and cytotoxic-effector programs. The concordant T-cell phenotypes observed in the mOE and hOE models further support the use of hOE tumors for subsequent EV-based pharmacologic studies (Figures 7A and S7F).

### NECTIN4 confers EV sensitivity and supports the activity of EV plus PD-1 blockade

Given that NECTIN4 determines EV targetability and is associated with both a CD8+ T-cell-inflamed microenvironment and higher ICI response rates in Ba/Sq tumors, we evaluated the activity of EV alone and in combination with PD-1 blockade.

In vitro, hOE in BOY-12E and MB49 cells substantially increased sensitivity to EV but not to monomethyl auristatin E (MMAE), the microtubule-disrupting cytotoxic payload of EV, consistent with NECTIN4-dependent EV activity rather than altered intrinsic sensitivity to MMAE (Figures S7J and S7K). Transcriptomic responses to EV and MMAE were highly concordant, supporting a dominant contribution of the MMAE payload to the transcriptional response to EV (Figures S7L and S7M).

In orthotopic models, EV significantly suppressed the growth of hOE tumors, whereas its activity was limited in EmV tumors (Figures 7I-7K and S7N). In hOE tumors, EV+anti-PD-1 antibody (EV+αPD-1) produced greater tumor suppression than EV alone, whereas neither EV nor the combination produced comparable tumor control in EmV tumors. Treatment was well tolerated with no substantial changes in body weight or peripheral blood counts (Figures S7O and S7P).

Given the preferential enrichment of TIGIT+CD8+ T cells in NECTIN4-high tumors, we next investigated whether TIGIT binding to tumor cells depended on NECTIN4 expression and whether EV altered this binding. Recombinant human TIGIT showed minimal binding to EmV cells but robust binding to hOE cells (Figure 8A). Notably, EV reduced TIGIT-Fc binding to hOE cells in a dose-dependent manner (Figures 8A and 8B), supporting the ability of EV to interfere with NECTIN4-dependent TIGIT binding. Together, these findings link NECTIN4 expression to EV targetability, a CD8-rich TME enriched for TIGIT-expressing T cells, and EV-sensitive TIGIT binding on the tumor cell surface. Thus, in addition to NECTIN4-dependent delivery of MMAE, interference with NECTIN4-dependent TIGIT binding may represent an additional immunomodulatory effect of EV.

**Figure 8.**
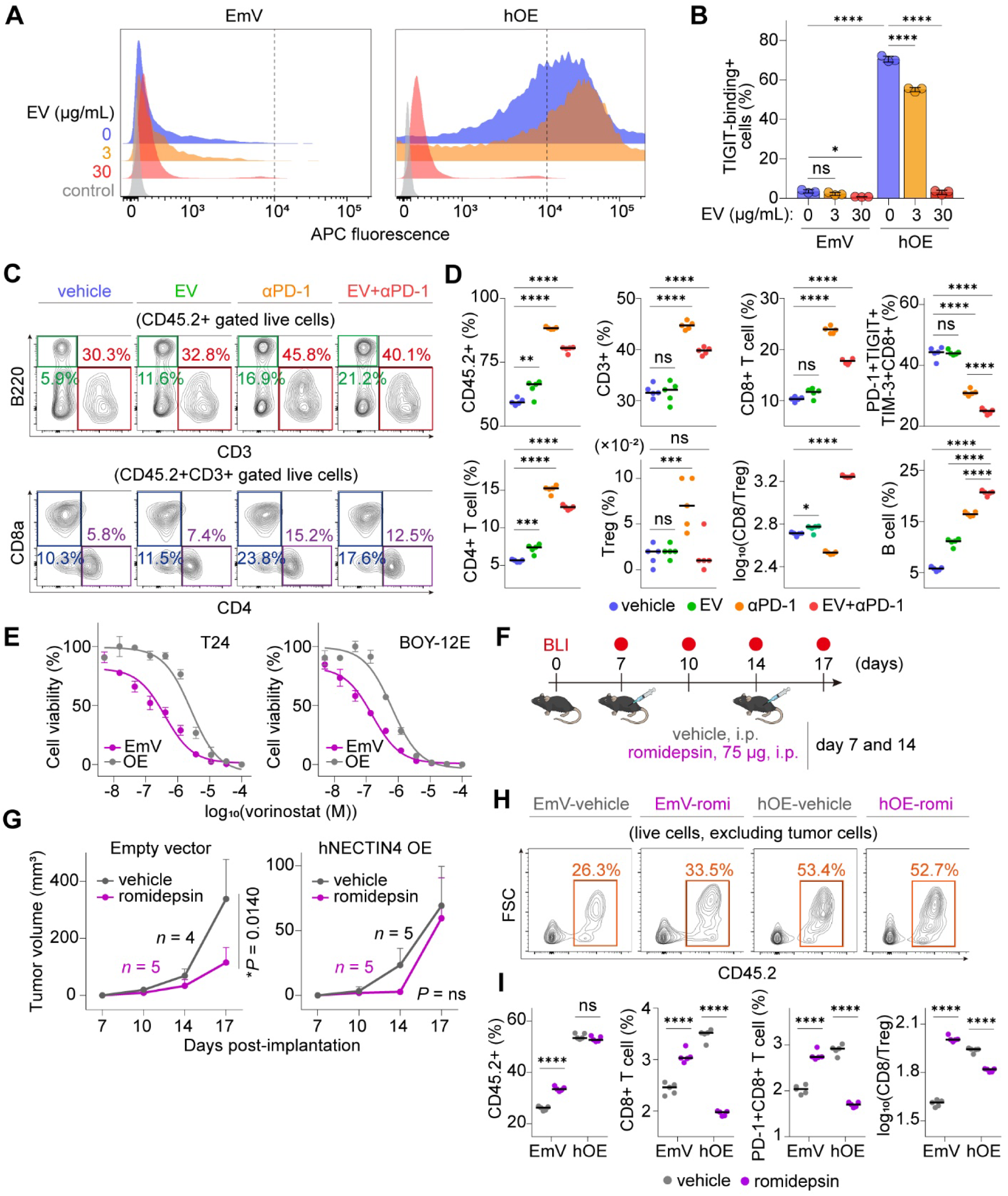
EV attenuates NECTIN4-dependent TIGIT binding and remodels the TME with PD-1 blockade, whereas NECTIN4-low models show preferential sensitivity to HDAC inhibition. (A-B) Recombinant TIGIT-Fc binding to empty-vector (EmV) and human NECTIN4-overexpressing (hOE) BOY-12E cells following enfortumab vedotin (EV) treatment, shown as histograms (A) and percentages of TIGIT-Fc+ cells (B). *n* = 3. (C-D) Flow-cytometric profiling (C) and frequencies of the indicated immune-cells and CD8/Treg ratios (D) in orthotopic hOE tumors treated with vehicle, EV, anti-PD-1 antibody (αPD-1), or EV+αPD-1. *n* = 5. (E) Dose-response curves for vorinostat in EmV and NECTIN4-overexpressing T24 and BOY-12E cells. *n* = 3. (F-I) Treatment schema (F), tumor growth (G) and immune profiling (H-I) of orthotopic EmV and hOE tumors following romidepsin treatment. *n* = 4 or 5. Data are mean ± SD in (B) and (E) and mean ± SEM in (G). Horizontal lines indicate medians. *P* values were determined by two-way ANOVA followed by Šídák’s multiple-comparisons test in and (I), one-way ANOVA followed by Tukey’s multiple-comparisons test in (D), and two-way ANOVA in (G). \**P* < 0.05, \*\**P* < 0.01, \*\*\**P* < 0.001 and \*\*\*\**P* < 0.0001; ns, not significant. BLI, bioluminescence imaging; romi, romidepsin; Treg, regulatory T cell.

### EV plus PD-1 blockade remodels T-cell and B-cell compartments in hOE tumors

Given the greater tumor suppression observed with EV+αPD-1 than with EV alone in hOE tumors, we next examined how combination treatment remodeled the TME. Orthotopic hOE tumors were treated with vehicle, EV, αPD-1, or EV+αPD-1, followed by multiparametric flow-cytometric profiling of tumor-associated immune cells (Figures S8A-S8D). αPD-1 and EV+αPD-1 were associated with higher frequencies of CD3+ T cells, including both CD8+ and CD4+ T-cell populations, than vehicle or EV treatment (Figures 8C and 8D). Notably, the frequency of PD-1+TIGIT+TIM-3+CD8+ T-cells was lower with EV+αPD-1 than αPD-1 monotherapy, indicating altered representation of a multi-checkpoint-expressing CD8+ T-cell compartment (Figure 8D). In contrast, αPD-1 monotherapy was associated with an increase in Treg frequency, which was attenuated in the EV and EV+αPD-1 groups; consequently, the combination group showed the highest CD8/Treg ratio (Figure 8D). These findings indicate that EV+αPD-1 reshapes the T-cell compartment toward increased CD8 relative to Treg representation.

EV+αPD-1 also increased B-cell frequency compared with either monotherapy (Figure 8D). The proportion of BCL6+ B cells was also higher in the combination group (Figure S8E), suggesting a treatment-associated shift in B-cell phenotype. Several myeloid populations also differed among treatment groups (Figure S8F). Overall, EV+αPD-1 induces a distinct immune composition characterized by increased T-cell representation and CD8/Treg ratio, reduced PD-1+TIGIT+TIM-3+CD8+ T cells relative to αPD-1 monotherapy, and increased B-cell frequency. These treatment-associated immune changes may accompany the greater tumor control observed with the combination in hOE tumors.

### HDAC inhibition preferentially targets the NECTIN4-low state

The limited activity of EV-based treatment in EmV tumors prompted us to ask whether the NECTIN4-low inflammatory-mesenchymal state harbors alternative therapeutic vulnerabilities. Given the inflammatory-mesenchymal programs associated with NECTIN4-low basal BC, and the reported ability of histone deacetylase inhibitors (HDACis) to modulate inflammatory cytokine signaling, including IL6-associated pathways^28–31^, we tested whether NECTIN4 status was associated with differential sensitivity to HDAC inhibition.

In vitro, EmV basal cells were consistently more sensitive to multiple HDACis than their hOE counterparts in both T24 and BOY-12E models (Figures 8E and S8G-S8H). These results identify preferential HDACi sensitivity as a reproducible feature of NECTIN4-low basal BC cell models.

We next evaluated romidepsin, a clinically approved HDACi^32^, in orthotopic tumors established with EmV or hOE cells (Figure 8F). Romidepsin significantly suppressed the growth of EmV tumors but had no significant effect on hOE tumor growth (Figure 8G). Treatment was well tolerated, with no substantial treatment-associated changes in the toxicity measures assessed (Figures S8I and S8J).

In EmV tumors, romidepsin increased the CD45.2+ leukocyte fraction, CD8+ T-cell representation, and the CD8/Treg ratio (Figures 8H-8I and S8K). By contrast, in hOE tumors, romidepsin reduced the CD8+ T-cell fraction, while DC and neutrophil fractions increased (Figures 8I and S8L). These findings show that HDAC inhibition has distinct antitumor and immune effects according to NECTIN4 expression in these models: romidepsin preferentially suppressed EmV tumor growth and was accompanied by increased leukocyte and CD8+ T-cell representation, whereas hOE tumors showed limited growth inhibition and a different pattern of immune-cell changes. Notably, the immune changes induced by romidepsin in EmV tumors— increased leukocyte infiltration, higher CD8+ T-cell representation, and an increased CD8/Treg ratio—mirror the baseline features that distinguish NECTIN4-high from NECTIN4-low tumors. This raises the possibility that HDAC inhibition partially converts the NECTIN4-low microenvironment toward the CD8-inflamed state associated with immunotherapy responsiveness, a hypothesis directly testable by combining HDAC inhibition with PD-1 blockade in this model.

## DISCUSSION

The remarkable clinical activity of EV+P has established NECTIN4 as a therapeutically important cell-surface target in urothelial carcinoma, yet the biology underlying this target remains incompletely understood. Here, we show that NECTIN4 is not simply an ADC antigen but a subtype-dependent regulator of tumor state and antitumor immunity in BC. The prognostic significance of NECTIN4 in BC has remained uncertain, with genomic and protein-level analyses yielding inconsistent associations with clinical outcomes^33–35^. Our subtype-stratified approach revealed that the association between high NECTIN4 expression and favorable clinical outcomes was largely restricted to Ba/Sq MIBC. Experimental perturbation further showed that NECTIN4 can restrain growth, migration, and inflammatory-mesenchymal programs in basal BC. This context-dependent activity contrasts with its established roles in other malignancies, where NECTIN4 has been reported to promote proliferation, metastasis, and EMT, including breast and pancreatic cancers^7,36–38^. Interestingly, this inter-tumor divergence extends beyond function to its baseline expression patterns: whereas NECTIN4 expression is generally low in basal BC, it is enriched in basal-like breast cancers^36,39^. This divergence may partly reflect tissue-specific transcriptional regulation of NECTIN4^40,41^. Together, these findings suggest that the biological consequences of NECTIN4 expression are strongly shaped by lineage and cellular context.

In BC, this context dependence may reflect distinct baseline transcriptional states across molecular subtypes. Ba/Sq MIBC is characterized by prominent inflammatory and mesenchymal programs, whereas LumP tumors generally exhibit less pronounced activity of these programs^10–12^. In basal BC models, NECTIN4 suppressed NFκB/IL6-STAT3-associated signaling and EMT-related transcriptional programs, and the reciprocal state was observed in human NECTIN4-low Ba/Sq tumors. Thus, NECTIN4-low Ba/Sq tumors appear to occupy an IL6/NFκB-high inflammatory-mesenchymal state associated with adverse clinical features. By contrast, NECTIN4 perturbation produced more limited growth and inflammatory-mesenchymal phenotypes in the luminal model, suggesting that the consequences of NECTIN4 expression depend on the underlying molecular context. This context dependence may contribute to the subtype-specific clinical associations observed in Ba/Sq MIBC.

An important implication of these findings is that tumor-associated inflammatory signaling and productive antitumor immune inflammation represent distinct biological states. Despite reduced IL6/NFκB and EMT-associated programs, NECTIN4-high Ba/Sq tumors exhibited a coordinated T cell-inflamed microenvironment characterized by increased CD8+ T-cell representation, spatial enrichment of immune cells around NECTIN4-positive tumor regions, and DC- and antigen-presentation-associated programs. In contrast, NECTIN4-low tumors combined inflammatory-mesenchymal programs with a less coordinated antitumor immune state.

Although Tregs are commonly associated with immunosuppression, their prognostic significance in BC has been inconsistent^42,43^. In our cohort, higher inferred Treg abundance was associated with favorable survival and co-occurred with increased CD8+ T-cell abundance, while CD8-high/Treg-high tumors showed favorable outcomes. These findings suggest that, in NECTIN4-high Ba/Sq tumors, Treg enrichment occurs within a broadly T cell-rich microenvironment rather than representing an isolated immunosuppressive feature.

Accordingly, TIGIT+CD8+ T cells enriched in NECTIN4-high tumors may represent checkpoint-expressing populations within an active T cell-inflamed state. This immune architecture is consistent with the higher ICI response rates observed in NECTIN4-high Ba/Sq tumors across two cohorts. In contrast, comparable NECTIN4-associated immune features were not observed in LumP tumors, consistent with the lack of a significant association between NECTIN4 status and ICI response in this subtype.

Intratumoral CD8+ T-cell infiltration is a well-established prognostic determinant in MIBC^44–46^, yet the tumor-intrinsic factors associated with this state remain incompletely understood. Our syngeneic mNectin4 model provides functional evidence that Nectin4 expression can promote CD8+ T-cell-dependent tumor control in vivo. Importantly, depletion of CD8+ T cells substantially diminished the tumor-control advantage conferred by mNectin4 expression, arguing that NECTIN4 is not merely a passive marker of a CD8-rich state. In the complementary hNECTIN4 model, immune-cell transcriptomes were enriched for antigen-processing machinery, MHC class I and II, DC-associated, interferon-response, and cytotoxic-effector programs. Together with the corresponding immune features observed in hNECTIN4-high tumors, these data support a link between the NECTIN4-high state and productive antitumor immunity.

The enrichment of TIGIT-expressing T cells in NECTIN4-high tumors raises an additional potential immunologic role for tumor-cell NECTIN4. Previous work has suggested that NECTIN4 can function as an inhibitory TIGIT ligand^47^, analogous to other NECTIN-family checkpoint interactions^48,49^. In line with this possibility, recombinant TIGIT preferentially bound NECTIN4-expressing BC cells, and EV reduced this binding in a dose-dependent manner. Although these experiments do not establish the functional consequences of NECTIN4-TIGIT engagement, they suggest that EV may interfere with TIGIT binding at the NECTIN4-expressing tumor-cell surface in addition to delivering its MMAE payload^15^. This raises the possibility that modulation of NECTIN4-dependent TIGIT binding contributes an additional immunologic component to EV activity, a hypothesis that will require direct functional testing.

EV plus PD-1 blockade was also associated with distinct remodeling of the immune compartment in hOE tumors. Compared with αPD-1 monotherapy, the combination reduced the relative representation of PD-1+TIGIT+TIM-3+CD8+ T cells and limited the increase in Tregs, resulting in the highest CD8/Treg ratio among the treatment groups. The combination also increased B-cell frequency and the proportion of BCL6+ B cells, indicating broader changes in the adaptive immune compartment. These findings suggest that the greater tumor control observed with EV plus PD-1 blockade is accompanied by immune remodeling beyond the direct cytotoxic activity of EV. Furthermore, EV-mediated bystander killing^50^ may additionally contribute to activity in tumors with spatially heterogeneous NECTIN4 expression.

A practical implication of this framework is that NECTIN4-low Ba/Sq disease may represent a biologically distinct therapeutic state characterized by reduced EV target availability and an IL6/NFκB-high inflammatory-mesenchymal program. Although our clinical cohorts did not receive EV+P and therefore cannot establish NECTIN4-low tumors as clinical EV+P nonresponders, NECTIN4-low basal BC models showed greater sensitivity to multiple HDAC inhibitors, and romidepsin preferentially suppressed NECTIN4-low tumors in vivo while increasing CD8+ T-cell representation and the CD8/Treg ratio. These findings nominate HDAC inhibition as an alternative vulnerability of the NECTIN4-low state and raise the possibility that epigenetic therapy may reshape this state toward a more immune-responsive phenotype. The molecular basis of this differential sensitivity and its potential interaction with checkpoint blockade remain to be determined. An early phase 1/1b study of the HDACi vorinostat plus pembrolizumab in advanced genitourinary cancers, including UC, demonstrated acceptable tolerability but limited antitumor activity in a biomarker-unselected population^51^. Our findings therefore provide a rationale for testing biomarker-informed approaches based on NECTIN4-defined tumor state.

In summary, our study identifies NECTIN4 not merely as a cell-surface target for ADCs but as a subtype-dependent regulator of tumor state and antitumor immunity in BC. In Ba/Sq tumors, NECTIN4 restrains NFκB/IL6-STAT3-associated inflammatory and mesenchymal programs while marking a spatially organized, CD8-rich immune state that is functionally linked to CD8+ T-cell-dependent tumor control. NECTIN4 thereby couples EV targetability to a subtype-specific tumor–immune context associated with checkpoint blockade responsiveness, whereas the NECTIN4-low state defines an inflammatory-mesenchymal phenotype with an alternative therapeutic vulnerability. More broadly, these findings establish a framework in which ADC targets can function not only as delivery antigens but also as biological determinants of tumor– immune state, with implications for understanding and rationally combining ADCs with immunotherapy.

## LIMITATIONS OF THE STUDY

Several limitations of this study warrant acknowledgment. First, the clinical analyses were retrospective, and selected validation cohorts, including the independent mIF and single-cell studies, had limited sample sizes. Moreover, because no patients received EV+P treatment in our cohort, prospective analyses of EV+P treated cohorts will be required to determine whether NECTIN4 expression and molecular subtype provide complementary predictive information. Second, the precise molecular mechanisms by which NECTIN4 regulates pro-tumorigenic inflammatory and EMT programs remain to be fully elucidated. Third, although EV reduced NECTIN4-dependent TIGIT binding, functional studies are needed to establish effects on CD8+ T cells and EV activity alone or with PD-1 blockade. Furthermore, while our mNectin4 models provided functional evidence for CD8-dependent tumor control, some immune-transcriptomic and EV-response experiments required expression of hNECTIN4 in murine MB49 cells. Although hOE tumors recapitulated key immune features observed in the mOE model and enabled pharmacologic evaluation of EV, a contribution from heterologous human NECTIN4 expression cannot be fully excluded. Moreover, both models rely on ectopic overexpression in cells with minimal endogenous Nectin4, and whether comparable effects occur at physiological endogenous expression levels remains to be determined.

## RESOURCE AVAILABILITY

### Lead contact

Further information and requests for resources and reagents should be directed to and will be fulfilled by the lead contact, Akihide Yoshimi.

### Material availability

Materials and reagents are available from the lead contact upon reasonable request. This study generated NECTIN4/Nectin4 expression constructs and stable cell lines, which are available from the lead contact upon reasonable request.

### Data and code availability

Raw and processed RNA-seq data generated from cell-line and mouse experiments have been deposited in the Gene Expression Omnibus (GEO) under accession number GSE349794. Bulk RNA-seq data from clinical samples, including processed expression matrices and deidentified sample metadata, have been deposited in the NCBI Gene Expression Omnibus under accession number GSE344756. Requests for access to additional deidentified data may be considered from qualified investigators upon execution of an appropriate data use agreement; requests should be directed to Akihide Yoshimi.

## ACKNOWLEDGMENTS

The authors thank Akiko Kagotani, Sayaka Sasada, and Rintaro Oide of the Translational Research Program, Osaka Medical and Pharmaceutical University, for their assistance with processing clinical samples used in this study. The authors also thank Junko Zenkoh of the Department of Computational Biology and Medical Sciences, Graduate School of Frontier Sciences, The University of Tokyo, for her assistance with the mIF experiments. The super-computing resource was provided by Human Genome Center (the University of Tokyo). This work was supported by the Grants-in-Aid for Scientific Research (A) (grant no. 21H04828 and 26H02447 to A.Y.) from the Ministry of Education, Culture, Sports, Science and Technology of Japan; the Research on Development of New Drugs from the Japan Agency for Medical Research and Development (AMED) (project number JP25ak0101263 to A.Y.); the Practical Research for Innovative Cancer Control from AMED (project number JP25ck0106906 to A.Y.); AMED-PRIME (project number JP24gm7010010 to A.Y.); and grants from Takeda Science Foundation and Canon Foundation (A.Y.).

## AUTHOR CONTRIBUTIONS

Conceptualization, K.Nishimura, K.K., and A.Y.; Methodology, K.Nishimura, T.N., K.K., S.K., and A.Y.; Validation, K.Nishimura, R.M., T.O., T.N., K.H., and S.K.; Formal analysis, K.Nishimura, T.N., K.H., and M.Ajiro; Investigation, K.Nishimura, R.M., T.O., M.Sakamoto, T.I.-T., M.Azuma, S.I., A.K., A.I., K.Nishimori, J.M., N.S., M.Sakumoto, M.Amisaki, K.O., M.I., Y.H., and Y.S.; Resources, K.Nishimura, R.M., T.T., Y.Y., H.A., K.K., F.L.F.C., A.S.K., L.J., and T.Y.; Data curation, K.Nishimura, R.M., K.H., T.T., and K.K.; Writing – original draft, K.Nishimura and A.Y.; Writing – review & editing, all authors; Visualization, K.Nishimura, T.N., and K.H.; Funding acquisition, K.K. and A.Y.; Supervision, H.N., H.A., S.K., K.K., and A.Y.

## DECLARATION OF GENERATIVE AI AND AI-ASSISTED TECHNOLOGIES IN THE MANUSCRIPT PREPARATION PROCESS

During the preparation of this work, the authors used ChatGPT (OpenAI) to improve the language and readability of the manuscript. After using this tool, the authors reviewed and edited the content as needed and take full responsibility for the content of the published article.

## STAR METHODS

### EXPERIMENTAL MODEL AND STUDY PARTICIPANT DETAILS

#### Clinical cohort and tissue specimens

Clinical and pathological data and biospecimens were retrospectively collected through the OMPU-NCC Consortium Japan, a collaborative bladder cancer (BC) consortium between Osaka Medical and Pharmaceutical University (OMPU) and the National Cancer Center Japan (NCC) ^52–55^. The study included 459 patients with histologically confirmed BC, comprising 143 patients with non-muscle-invasive bladder cancer (NMIBC) and 316 patients with muscle-invasive bladder cancer (MIBC), as well as 35 histologically normal tumor-adjacent bladder tissue samples.

Tumor specimens were obtained by transurethral resection of bladder tumor or radical cystectomy at OMPU. Fresh tumor and tumor-adjacent normal bladder tissues were collected for molecular analyses, whereas corresponding formalin-fixed paraffin-embedded (FFPE) specimens were used for tissue microarray (TMA) construction, immunohistochemistry (IHC), and spatial immune profiling. Hematoxylin and eosin (H&E)-stained sections were reviewed by board-certified pathologists to confirm the pathological diagnosis and verify the absence of tumor cells in tumor-adjacent normal tissues. Detailed procedures for tissue preservation, molecular quality control, nucleic acid extraction, and sequencing are described below.

The study was conducted in accordance with the Declaration of Helsinki and applicable regulations governing noninterventional human studies. The collection and use of clinical information and human biospecimens were approved by the Institutional Review Boards of OMPU (approval numbers 2305-11, 2344-10, 2523-3, 2808-2, and 2571-6) and NCC (approval numbers 2020-486, 2022-125, 2023-290, and 2024-232). Written informed consent was obtained from all participants before the use of their clinical information and biospecimens.

### Animal experiments

MB49 cells were transduced with the pHAGE PGK-GFP-IRES-LUC-W plasmid (Addgene, plasmid #46793), and green fluorescent protein (GFP)-positive cells were isolated using a FACSAria III cell sorter (BD Biosciences). Female C57BL/6 mice, 6-8 weeks of age, were purchased from CLEA Japan, Inc. (Tokyo, Japan). Mice were anesthetized with isoflurane, and 5 × 10^4^ MB49 cells suspended in 10 μL Matrigel were orthotopically injected into the bladder dome using a Hamilton microsyringe equipped with a 30-gauge needle. Four days after tumor implantation, tumor establishment was confirmed by bioluminescence imaging (BLI) using an IVIS Lumina LT system (PerkinElmer), and tumor-bearing mice were then randomized into the indicated treatment groups. No statistical method was used to predetermine sample size. No animals were excluded after randomization unless prespecified humane endpoints, such as excessive body-weight loss or signs of severe distress, were met. Investigators performing BLI quantification were blinded to treatment group. Tumor growth was monitored by BLI, and bioluminescence intensity was quantified as radiance (photons/sec/cm²/sr). Tumor size was monitored every 3-4 days by BLI using the IVIS system.

For combination-treatment experiments, mice received vehicle, enfortumab vedotin (EV) (15 μg per mouse), an anti-mouse PD-1 antibody (150 μg per mouse), or their combination by intraperitoneal (i.p.) injection on days 7, 14 and 21 after tumor implantation. Tumor growth was monitored by BLI on days 7, 10, 14, 17 and 21. For CD8⁺ T-cell depletion experiments, mice received an anti-mouse CD8β antibody or the corresponding IgG1 isotype control at 50 μg per mouse i.p. on days -1, 0, 7 and 14 relative to tumor implantation. For histone deacetylase (HDAC) inhibition experiments, romidepsin was administered i.p. at 75 μg per mouse on days 7 and 14 for tumor-growth studies and on days 7, 14 and 21 for immune-profiling studies. Tumors in the latter experiments were collected on day 26 for flow-cytometric analysis.

Body weight and general condition were monitored throughout the experiments. At the indicated endpoints, mice were euthanized, and bladder tumors and peripheral blood were collected for further analyses. All animal experiments were approved by the Animal Care and Use Committee of the National Cancer Center under protocol number T21-003-M18.

#### Cell lines

T24, 5637, HT-1376 and MCF7 cells were obtained from the American Type Culture Collection (ATCC); BOY-12E cells from the RIKEN BioResource Research Center; KMBC-2 cells from the JCRB Cell Bank; MB49 cells from CLS Cell Lines Service GmbH; and Lenti-X 293T cells from Takara Bio. T24, 5637, and BOY-12E cells were cultured in RPMI 1640; KMBC-2 and MB49 cells were cultured in Dulbecco’s modified Eagle medium (DMEM); HT-1376 and MCF7 cells were cultured in Eagle’s Minimum Essential Medium; and Lenti-X 293T cells were cultured in high-glucose DMEM. All media were supplemented with 10% fetal bovine serum and 1% penicillin-streptomycin. Cells were maintained at 37°C in a humidified atmosphere containing 5% CO_2_. Cell lines were obtained directly from the indicated repositories or vendors. No additional cell line authentication was performed in-house. All cell lines were routinely screened and confirmed to be free of mycoplasma contamination.

### METHOD DETAILS

#### NECTIN4 IHC

FFPE TMA sections were stained with an anti-human NECTIN4 antibody (clone EPR15613-68; Abcam, ab192033; 1:1,000) using the Discovery ULTRA automated staining system (Roche Diagnostics, Basel, Switzerland) according to the manufacturer’s protocol. NECTIN4 staining was independently evaluated by at least two board-certified pathologists who were blinded to clinical outcomes. Discrepant assessments were resolved by consensus. NECTIN4 expression was quantified using an H-score calculated as (% cells with weak staining × 1) + (% cells with moderate staining × 2) + (% cells with strong staining × 3), yielding a score from 0 to 300. An H-score of 100, corresponding to the prespecified boundary between low (15–99) and moderate (100–199) staining categories, was used to classify tumors as NECTIN4-low or NECTIN4-high.

#### Nucleic acid extraction, whole-exome sequencing, and RNA sequencing (RNA-seq) of clinical specimens

Fresh tumor and tumor-adjacent normal bladder tissues were immediately preserved in RNAlater (Thermo Fisher Scientific). For molecular profiling, H&E-stained sections were reviewed by board-certified pathologists, and tumor specimens containing at least 60% tumor-cell nuclei and no more than 20% necrotic tissue were included. DNA and RNA were extracted using the AllPrep DNA/RNA Kit (QIAGEN) according to the manufacturer’s instructions. Nucleic acid concentrations were measured using a NanoDrop Microvolume UV-Vis Spectrophotometer (Thermo Fisher Scientific). RNA integrity was assessed using an Agilent 2100 Bioanalyzer (Agilent), and samples with an RNA integrity number below 7.0 were excluded from RNA-seq analysis.

Library preparation was automated using Maholo LabDroids at the Robotic Biology Institute Inc. (Tokyo, Japan). For RNA-seq, ribosomal RNA was depleted using the NEBNext rRNA Depletion Kit, followed by strand-specific library preparation using the NEBNext Ultra II Directional RNA Library Prep Kit for Illumina (New England Biolabs) according to the manufacturer’s instructions.

For whole-exome sequencing, exonic regions from tumor and matched normal DNA were captured using the xGen Exome Research Panel v1.0 (Integrated DNA Technologies), and libraries were prepared using the KAPA HyperPlus Kit (KAPA Biosystems) according to the manufacturer’s instructions. Whole-exome and RNA-seq libraries were sequenced on an Illumina NovaSeq 6000 platform. Whole-exome sequencing was performed using 150-bp paired-end reads with a target mean coverage of at least 200×, whereas RNA-seq was performed using 100-bp paired-end reads with a target of at least 50 million reads per sample.

#### RNA extraction and RNA-seq of cultured cells

Total RNA was extracted from T24 and BOY-12E cells stably overexpressing human NECTIN4 (OE) or the corresponding empty-vector (EmV) control, and from KMBC-2 cells with stable NECTIN4 knockdown or the corresponding control. For drug-treatment experiments, BOY-12E cells were treated for 4 days with EV or monomethyl auristatin E (MMAE) at concentrations that produced approximately 20% growth inhibition, or with vehicle control, before RNA extraction.

Total RNA was isolated using the FavorPrep Tissue Total RNA Kit (FAVORGEN), followed by DNase I treatment (QIAGEN), according to the manufacturers’ instructions. Strand-specific mRNA libraries were prepared and sequenced either by BGI Tech Solutions using the DNBSEQ Eukaryotic Strand-specific Transcriptome Resequencing service or by Novogene Japan Co., Ltd. using its directional RNA-seq service, according to the respective service providers’ standard procedures.

#### RNA-seq of tumor-infiltrating CD45.2⁺ cells

Orthotopic tumors generated from MB49 EmV or human NECTIN4-OE cells were dissociated into single-cell suspensions. CD45.2+ immune cells were isolated using a FACSAria III cell sorter (BD Biosciences). Total RNA was extracted from the sorted cells using the FavorPrep Tissue Total RNA Extraction Mini Kit (FAVORGEN), followed by DNase I treatment (QIAGEN). Strand-specific RNA-seq libraries were prepared and sequenced by Novogene Japan Co., Ltd. using its directional RNA-seq service according to the service provider’s standard procedures.

#### Bioinformatic analyses

Whole-exome and RNA-seq data from clinical specimens were processed using the SHIROKANE supercomputing system at the Human Genome Center, Institute of Medical Science, The University of Tokyo. Processing of whole-exome sequencing data, including sequence alignment, somatic variant calling, and copy-number analysis, was performed as previously described^52^.

For RNA-seq data from the OMPU-NCC cohort, sequencing reads were aligned to the human reference genome GRCh38 using STAR (v2.7.11b)^56^ with gene annotations from Ensembl release 112. Gene-level read counts were quantified using featureCounts (v2.0.6)^57^ and converted to transcripts per million (TPM) values. To quantify pathway activity, a z-score was calculated for each gene across all samples, and pathway-level scores were calculated by averaging the z-scores of genes included in each Molecular Signatures Database (MSigDB) Hallmark gene set.

Differential gene expression analysis of clinical specimens was performed using raw gene-count matrices using DESeq2 implemented in iDEP v1.13^58^. Principal component analysis was performed using the same platform. Differentially expressed genes were defined using |log2 fold change| > 1 and adjusted P < 0.05. Pathway enrichment analysis of differentially expressed gene lists was performed using ShinyGO v0.80^59^.

For RNA-seq data from cultured cells and sorted CD45.2+ cells, sequence reads were aligned to the GRCh38 or GRCm39 reference genome using STAR (v2.7.10b)^56^ and quantified at the gene level using StringTie2 (v2.2.1)^60^ and converted to TPM values. Differential gene expression and principal component analyses were performed using DESeq2 implemented in iDEP v1.13^58^. Differentially expressed genes were defined using |log2 fold change| > 1 and adjusted P < 0.05. Pathway enrichment analysis of differentially expressed gene lists was performed using ShinyGO v0.80^59^. Gene set enrichment analysis (GSEA) was performed using the Broad Institute GSEA desktop application with the MSigDB Hallmark gene sets, using 1,000 gene-set permutations, the signal-to-noise ratio for gene ranking, and a weighted enrichment statistic. Protein-protein interaction networks were generated using the STRING database^61^ and visualized using Cytoscape software^62^. Bubble plots were generated using the SRplot web platform^63^.

#### Molecular subtype classification

MIBC molecular subtypes were assigned from bulk RNA-seq data using the consensusMIBC R package (version 1.1.0) according to the consensus molecular classification^12^. NMIBC tumors were assigned to the UROMOL 2021 system using the classifyNMIBC R package (version 1.1.0)^13^. The same consensusMIBC classifier was applied to the IMvigor210 dataset.

#### CIBERSORTx and EcoTyper analyses

TPM-normalized gene expression matrices generated as described above were used for CIBERSORTx and EcoTyper analyses. CIBERSORTx^64^ was performed in absolute mode using the LM22 reference signature matrix to estimate the abundance of 22 immune-cell subsets. EcoTyper^23^ was applied using the predefined carcinoma model to assign transcriptional profiles to previously characterized cell states and ecotypes.

#### Immune-related gene signature scoring

For RNA-seq data from tumor-infiltrating CD45.2⁺ cells, antigen-processing machinery/major histocompatibility complex class I (APM/MHC-I), MHC class II (MHC-II), and dendritic-cell (DC) priming scores were calculated as the mean of the gene-wise z-scores for genes included in each signature. The APM/MHC-I signature comprised H2-K1, H2-D1, B2m, Tap1, Tap2, Tapbp, Psmb8, Psmb9, Psmb10, and Erap1. The MHC-II signature comprised Ciita, Cd74, H2-Aa, H2-Ab1, H2-Eb1, H2-Dma, H2-DMb1, and Ctss. The DC priming signature comprised Cd80, Cd86, Cd40, and Il12b.

Cytolytic activity (CYT) was calculated as the geometric mean of Gzma and Prf1 expression, as previously described^65^, with a pseudocount of 0.01 added to each TPM value, as follows: CYT = √[(Gzma TPM + 0.01) × (Prf1 TPM + 0.01)]. CYT scores were subsequently log2-transformed for downstream analysis and visualization.

#### Analysis of the IMvigor210 and The Cancer Genome Atlas bladder cancer cohorts

Publicly available RNA-seq and clinical data from the IMvigor210 trial of atezolizumab in patients with metastatic urothelial carcinoma (UC) were obtained from the publicly released IMvigor210 dataset^14^. Patients were classified as NECTIN4-high or NECTIN4-low using the median NECTIN4 expression level as the cutoff.

To assess the frequency of NECTIN4 copy-number amplification in BC, putative copy-number alteration data from the Bladder Urothelial Carcinoma cohort of The Cancer Genome Atlas (TCGA-BLCA) PanCancer Atlas^11^ were queried using cBioPortal^66^.

#### CODEX/PhenoCycler spatial imaging and analysis

FFPE TMA sections were stained using the PhenoCycler platform (Akoya Biosciences) with a 41-marker DNA-barcoded antibody panel. Iterative annealing of fluorescent oligonucleotide reporters, image acquisition, and reporter stripping were performed according to the FFPE-optimized CODEX workflow. Image registration and single-cell signal extraction were subsequently performed.

Individual cells were segmented using QuPath software (v0.4.4)^67^ based on nuclear masks together with membrane and cytoplasmic markers. Cell phenotypes were assigned according to predefined marker combinations, and per-cell marker intensities, phenotype labels, and x-y spatial coordinates were exported as flow cytometry standard (FCS) and comma-separated values (CSV) files for downstream analysis. Detailed antibody panels and phenotype definitions are provided in Table S3.

Spatial analyses were performed using CytoMAP in MATLAB^68^. Single-cell coordinates and phenotype assignments were imported, and tissues were subdivided into evenly spaced, raster-scanned circular neighborhoods with a radius of 50 μm and a grid step of 25 μm. Local cellular composition was quantified within each neighborhood. Cellular neighborhoods were operationally defined as groups of spatial neighborhoods with similar cellular compositions. Neighborhoods were clustered into spatial tissue regions using k-means clustering, with the optimal number of regions determined using the Davies-Bouldin criterion. Region assignments were mapped back onto the original tissue coordinates for visualization. Neighborhood-level correlation and nearest-neighbor analyses were performed to evaluate spatial associations between NECTIN4-positive tumor cells and immune-cell populations.

#### Multiplex Immunofluorescence (mIF)

After deparaffinization, antigen retrieval (AR) was performed using a heat-induced epitope retrieval method with 10X AR9 Buffer (Akoya Bioscience). The tissue sections were blocked with Antibody Diluent with Background-Reducing Components (Agilent) for 10 minutes, followed by a 30-minute incubation with a primary antibody. Antibody dilutions were as follows: PD-1, 1:200; FOXP3, 1:400; TIGIT, 1:400; CD8, 1:100; CD3, 1:200; pan-cytokeratin, 1:400; HLA-DR, 1:1,000; CD163, 1:500; CD11c, 1:100; CD66b, 1:100; and CD68, 1:200. Afterward, the sections were incubated with an Opal Polymer HRP secondary antibody (Akoya Biosciences) and stained with Opal fluorophores (Opal 480) at a dilution of 1:100 for 10 minutes. AR was repeated using 10X AR6 Buffer (Akoya Bioscience) to prepare the slides for subsequent antibody application. After the same steps were repeated from blocking to Opal reagent application (Opal 480, 520, 570, 620, 690, and 780) at a dilution of 1:100, the tissue slides were counterstained with 10X Spectral DAPI (Akoya Bioscience) for nuclear visualization for 10 minutes and mounted with coverslips.

The stained slides were scanned using PhenoImager Fusion (Akoya Biosciences). A supervised machine-learning classifier was trained using manually annotated representative areas to distinguish pan-cytokeratin-positive tumor nests from pan-cytokeratin-negative tumor stroma. Tumor sections were randomly selected and annotated using inForm analysis software (Akoya Biosciences). Cell count and definition of immune cell were automatically quantified using HALO v3.6 (Indica Labs). Three representative regions of interest from each compartment were selected for each case by a pathologist who was blinded to the clinical information and outcomes. CD8+ T cells were defined as CD3+CD8+FOXP3-cells, and regulatory T cells (Tregs) were defined as CD3+FOXP3+CD8-cells. PD-1+ and TIGIT+CD8+ T cells were defined as CD3+CD8+FOXP3-cells expressing PD-1 or TIGIT, respectively. Neutrophils were defined as CD66b+ cells, and dendritic cells were defined as CD11c+HLA-DR+ cells. Macrophages were identified by CD68 expression and operationally classified as M1-like macrophages (CD68+HLA-DR+CD163-) or M2-like macrophages (CD68+CD163+HLA-DR-). The density of each immune-cell population was calculated as the number of phenotypically defined cells per mm² and quantified separately within the tumor-nest and stromal compartments.

#### Analysis of publicly available single-cell RNA-seq (scRNA-seq) datasets

Publicly available scRNA-seq datasets of MIBC tumors were obtained from the Gene Expression Omnibus (GEO; GSE186520) for the Wu et al. cohort^27^ and from Mendeley Data for the Salomé et al. cohort (DOI: 10.17632/7yb7s9769c.1)^26^. Gene expression profiles were aggregated at the sample level to generate pseudo-bulk transcriptomes and converted to TPM values. Molecular subtypes were assigned using the consensusMIBC package (v1.1.0)^12^. NECTIN4-high and -low tumors were defined using a pseudo-bulk NECTIN4 expression cutoff of 5 TPM.

Cells with fewer than 500 unique molecular identifier counts were excluded, and doublets were identified and removed using scDblFinder (v1.8.0)^69^. Additional low-quality cells were filtered using sample-specific thresholds for the number of detected genes and mitochondrial transcript abundance. Downstream analyses were performed using Seurat (v5.5.0)^70^. Data were log-normalized with a scale factor of 10,000, and 5,000 highly variable genes were identified. Principal component analysis was performed, followed by batch correction and dataset integration using Harmony (v1.2.3)^71^. The first 30 Harmony-adjusted principal components were used for nearest-neighbor graph construction and clustering.

T-cell clusters were identified based on canonical marker expression and reclustered using the first 10 Harmony-adjusted principal components. Because the proliferating T-cell cluster contained both lineages, this cluster was further subclustered using the first 5 Harmony-adjusted principal components, and the resulting CD4+ and CD8+ subclusters were merged with the corresponding T-cell populations. Cells with discordant lineage assignments were manually reassigned based on CD4 and CD8B expression using a threshold of 0.5, whereas cells expressing both markers above this threshold were considered ambiguous and excluded.

CD4+ and CD8+ T cells were subsequently projected onto reference tumor-infiltrating lymphocyte (TIL) atlases using ProjecTILs (v3.7.0)^72^. The human CD4+ TIL reference atlas derived from Zheng et al.^73^ and the human CD8+ TIL reference atlas compiled from publicly available datasets, as listed in the Key Resources Table, were used for fine T-cell state annotation. Cells with log-normalized TIGIT expression >1.5 were defined as TIGIT-high, using a selected threshold corresponding approximately to the first quartile of TIGIT expression among TIGIT-expressing cells after excluding cells with near-zero expression.

#### Generation of NECTIN4/Nectin4 expression plasmids and lentiviral transduction

The coding sequence of human NECTIN4 (NCBI Reference Sequence NM_030916.3) was amplified from MCF7-derived complementary DNA (cDNA) using the following primers: forward, 5′-AAAAGGATCCACCATGCCCCTGTCCCTG-3′; and reverse, 5′-AAAAGCTAGCTGGGTCAGACCAGGTGTCC-3′. A pLX304-derived lentiviral backbone was generated from V5-APEX2-SENP2 (Addgene, #129276) by removing the entire V5-APEX2-SENP2 expression cassette. The backbone was amplified using the following primers: forward, 5′-AAAAGGATCCTTCTATCTCCTTCGAAGCCTGC-3′; and reverse, 5′-CCGAGCTTGGGTTTGCTGAT-3′. The human NECTIN4 insert was cloned into the modified pLX304 backbone using the BamHI and NheI restriction sites to generate pLX304-hNECTIN4.

A matched EmV control lacking the NECTIN4 coding sequence was generated in parallel. The coding sequence of mouse Nectin4 (NCBI Reference Sequence NM_027893.3) was synthesized and cloned into the same modified pLX304 backbone by GenScript to generate pLX304-mNectin4. The integrity of the resulting constructs was confirmed by Sanger sequencing.

Lentiviral particles encoding human NECTIN4, mouse Nectin4, or the corresponding EmV control were produced by transient transfection of Lenti-X 293T packaging cells. Cells cultured in 10-cm dishes were transfected with 10 μg of pLX304-hNECTIN4, pLX304-mNectin4, or the corresponding EmV plasmid, together with 4 μg of pCMV-VSV-G (Addgene, #8454) and 6 μg of psPAX2 (Addgene, #12260), using 70 μL of PEI MAX solution (pH 7.0) in 500 μL of Opti-MEM (Thermo Fisher Scientific). The culture medium was replaced 6 h after transfection. Viral supernatants were harvested 48 h after transfection and clarified by centrifugation at 4,000 rpm for 5 min to remove cellular debris. BOY-12E, T24, and MB49 cells were exposed to the clarified viral supernatants for 48 h. Transduced cells were subsequently selected with blasticidin at 10 μg/mL for 7 days.

#### Generation of NECTIN4-knockdown cells

Short hairpin RNAs (shRNAs) targeting human NECTIN4 were cloned into the AgeI and EcoRI sites of pLKO.1-puro (Addgene, #8453). The empty pLKO.1-puro vector was used as the control. The shRNA target sequences were as follows: shNECTIN4-1, 5′-CCCTTTCCATTAGTTGTATTT-3′; and shNECTIN4-2, 5′-CCAGCAGATGACCCAGAAATA-3′. Lentiviral particles were produced and transduced into target cells as described above. Transduced cells were selected with puromycin at 5 μg/mL for 7 days. For RNA-seq analysis, KMBC-2 cells expressing shNECTIN4-1 and the corresponding empty-vector control cells were used.

#### Colony formation assay

A total of 600 cells per well were seeded in six-well plates and cultured at 37°C for 10 days. Colonies were fixed with 4% paraformaldehyde (FUJIFILM Wako Pure Chemical Corporation) for 30 min and stained with 0.01% crystal violet (FUJIFILM Wako Pure Chemical Corporation) for 30 min. Colonies greater than 1 mm in diameter were independently counted by two researchers. All experiments were performed in triplicate.

#### Scratch migration assay

A total of 6 × 10^5^ cells per well were seeded in six-well plates and cultured until they reached 100% confluence. A linear scratch was created across the center of each well using a 200-μL pipette tip. The cells were incubated at 37°C, and images of the scratched areas were acquired every 6 hours using an EVOS imaging system (Thermo Fisher Scientific). Wound area was quantified using ImageJ, and migration was expressed as percentage wound closure relative to the area at 0 h. All experiments were performed in triplicate.

#### Semi-quantitative reverse transcription PCR (RT-PCR) and quantitative real-time PCR (qPCR)

Primer sequences used for semi-quantitative RT-PCR and qPCR were as follows: human NECTIN4, forward 5′-CAGTTACTCCACGCTGACCA-3′ and reverse 5′-TGTTTGATGCCTTCATCCTG-3′; human GAPDH, forward 5′-GCAAATTCCATGGCACCGTC-3′ and reverse 5′-CCAAAATCAAGTGGGGCGA-3′; and mouse Gapdh, forward 5′-GGTGCTGAGTATGTCGTGGA-3′ and reverse 5′-CACACCCATCACAAACATGG-3′.

cDNA was synthesized from total RNA using ReverTra Ace qPCR RT Master Mix (TOYOBO). For semi-quantitative RT-PCR, amplified products were separated by agarose gel electrophoresis and visualized using an iBright Imaging System (Thermo Fisher Scientific). qPCR was performed using SYBR Green Master Mix (Roche) on a CFX Duet Real-Time PCR System (Bio-Rad). Each sample was analyzed in technical triplicate. Target-gene expression was normalized to the corresponding GAPDH/Gapdh expression and quantified using the comparative Ct method.

#### Immunoblotting

Whole-cell lysates were prepared in radioimmunoprecipitation assay (RIPA) buffer (FUJIFILM Wako Pure Chemical Corporation) supplemented with protease inhibitor cocktail (Thermo Fisher Scientific). Lysates were clarified by centrifugation at 13,500 rpm for 5 min. Protein concentrations were determined using the Bradford Protein Assay (Bio-Rad), and equal amounts of protein were separated by sodium dodecyl sulfate-polyacrylamide gel electrophoresis (SDS-PAGE) using 4%-20% Mini-PROTEAN TGX precast gels (Bio-Rad) and transferred onto polyvinylidene difluoride (PVDF) membranes. Membranes were blocked with 5% bovine serum albumin in Tris-buffered saline containing 0.1% Tween 20 (TBST) for 1 h at room temperature and incubated overnight at 4°C with the primary antibodies listed in the Key Resources Table. After incubation with the corresponding horseradish peroxidase (HRP)-conjugated secondary antibodies, signals were detected using an enhanced chemiluminescence reagent (Cytiva) and imaged using an iBright CL1500 Imaging System (Thermo Fisher Scientific). When necessary, membranes were stripped using Stripping Solution (FUJIFILM Wako Pure Chemical Corporation) according to the manufacturer’s instructions and reprobed with the indicated antibodies. Immunoblot band intensities were quantified using ImageJ/Fiji. Phosphorylated protein levels were normalized to the corresponding total protein levels. Values were then normalized to the corresponding EmV control.

#### Immunofluorescence

T24 and BOY-12E EmV and NECTIN4-OE cells were seeded onto poly-D-lysine-coated glass coverslips. Cells were fixed with 4% paraformaldehyde in phosphate-buffered saline (PBS) for 15 min, permeabilized with 0.1% Triton X-100 in PBS for 15 min, and blocked with 3% bovine serum albumin in PBS for 1 h at room temperature. Cells were incubated overnight at 4°C with antibodies against NECTIN4 (1:200) or phospho-NF-κB p65 (1:800). After washing with PBS containing 0.1% Tween 20 (PBST), cells were incubated with the corresponding conjugated secondary antibodies (1:1000) for 1 h at room temperature. Nuclei were counterstained with Hoechst 33342. Images were acquired using a FLUOVIEW FV3000 Confocal Laser Scanning Microscope (Olympus/Evident) or a CKX53 inverted fluorescence microscope (Olympus/Evident), as appropriate.

#### Cell viability assay

A total of 3,000 cells per well were seeded in 96-well plates and incubated overnight at 37°C. Cells were treated with vehicle, MMAE, EV, entinostat, trichostatin A (TSA), romidepsin, or vorinostat for 4 days without medium replacement. Cell viability was measured using the CellTiter-Glo 2.0 Cell Viability Assay (Promega) according to the manufacturer’s instructions. All conditions were tested in triplicate, and dose–response curves were generated to assess drug sensitivity.

#### Recombinant TIGIT-binding assay

Recombinant human TIGIT-Fc chimera protein (5 µg/mL; R&D Systems) and human IgG1 isotype control (Selleck) were used as primary probes. BOY-12E EmV and NECTIN4-OE cells were suspended at 1 × 10^5^ cells per 100 µL in staining buffer consisting of PBS supplemented with 5% bovine serum albumin and 0.05% sodium azide. Cells were preincubated with EV at the indicated concentrations on ice for 60 min. Recombinant human TIGIT-Fc or the corresponding human IgG1 isotype control was then added, and cells were incubated on ice for 1 h. After thorough washing with staining buffer, cells were incubated with allophycocyanin (APC)-conjugated AffiniPure F(ab′)_2_ fragment donkey anti-human IgG, Fcγ fragment-specific antibody (Jackson ImmunoResearch) on ice for 30 min. Cells were subsequently washed three times, resuspended in FACFlow containing DAPI, and analyzed using a FACSAria III flow cytometer (BD Biosciences).

TIGIT-Fc binding was quantified after gating on singlets and excluding DAPI-positive dead cells. Ethanol-treated cells were used as a positive control to establish the dead-cell gate. To control for nonspecific IgG and secondary-antibody binding as well as APC fluorescence arising from cell-associated EV, cells were preincubated with 30 µg/mL EV and incubated with the human IgG1 isotype control, followed by the APC-conjugated secondary antibody. Data were analyzed using FlowJo v10. TIGIT-Fc-positive cells were defined as cells with APC fluorescence intensity ≥10^4^.

#### Tumor dissociation and flow cytometry

At the indicated experimental endpoints, mice were euthanized and bladder tumors were collected for flow-cytometric analysis. Tumors were minced into small pieces and incubated in 5 mL of digestion medium containing 500 μL of collagenase/hyaluronidase, 750 μL of DNase I, and 3.75 mL of RPMI 1640 to generate single-cell suspensions, following the previously described tumor-dissociation procedure.

Single-cell suspensions were stained with Zombie Aqua Fixable Viability Dye (BioLegend) and fluorochrome-conjugated anti-mouse antibodies against CD45.2, CD3, CD4, CD8α, CD25, PD-1, TIGIT, TIM-3, B220, CD11b, CD11c, Ly6G, F4/80, and CD163. Intracellular staining for FOXP3 and BCL6 was performed after fixation and permeabilization using a Fixation/Permeabilization buffer (eBioscience) according to the manufacturer’s instructions. After gating on singlets and Zombie Aqua-negative viable cells, GFP-positive tumor cells were excluded, and CD45.2-positive leukocytes and their respective immune-cell subsets were analyzed. Data were acquired using a FACSAria III flow cytometer (BD Biosciences) and analyzed using FlowJo v10 (BD Biosciences).

T cells were defined as CD45.2+CD3+ cells; CD8+ T cells as CD45.2+CD3+CD8α+CD4-cells; CD4+ T cells as CD45.2+CD3+CD4+CD8α-cells; B cells as CD45.2+B220+CD3-cells; and Tregs as CD45.2+CD3+CD4+CD25+FOXP3+ cells. PD-1+CD8+ T cells and PD-1+TIGIT+CD8+ T cells, and PD-1+TIGIT+TIM-3+CD8+ T-cell subsets were defined according to expression of the indicated markers within CD45.2+CD3+CD8α+ T cells. Neutrophils were defined as CD45.2+CD3-Ly6G+ cells; macrophages as CD45.2+CD3-Ly6G-CD11b+F4/80+ cells; CD163+ M2 macrophages as CD45.2+CD3-Ly6G-CD11b+F4/80+CD163+ cells; and dendritic cells as CD45.2+CD3-Ly6G-CD11c+F4/80-cells.

### QUANTIFICATION AND STATISTICAL ANALYSIS

#### Clinical statistical analysis

Overall survival was defined as the interval from tumor resection to death from any cause or the last follow-up. Progression-free survival was defined as the interval from tumor resection to radiographically or clinically confirmed disease progression. Cancer-specific survival was defined as the interval from tumor resection to death attributable to BC or the last follow-up. Responses to immune checkpoint inhibitors were evaluated according to Response Evaluation Criteria in Solid Tumors version 1.1. Complete and partial responses were classified as responses, whereas stable and progressive disease were classified as nonresponses.

Categorical variables were compared using Fisher’s exact test or Pearson’s chi-square test. The distribution of continuous variables was evaluated using the Kolmogorov-Smirnov test. Normally distributed variables were compared using Student’s t test or one-way analysis of variance (ANOVA), whereas non-normally distributed variables were compared using the Mann-Whitney U test or Kruskal-Wallis test. Survival curves were estimated using the Kaplan-Meier method and compared using the log-rank test. Univariable Cox proportional hazards regression was performed to estimate hazard ratios and 95% confidence intervals for overall survival. A multivariable Cox proportional hazards model was subsequently fitted including NECTIN4 expression, clinical stage, prior radiotherapy, prior chemotherapy, prior BCG treatment, lymphovascular invasion, sex, and age. The proportional hazards assumption was assessed using Schoenfeld residuals. Associations between continuous variables were evaluated using simple linear regression, as appropriate. All statistical tests were two-sided, and *P* < 0.05 was considered statistically significant. Clinical statistical analyses were performed using GraphPad Prism version 9.3.1, JMP Pro version 17, and R version 4.6.1. Kaplan-Meier curves and forest plots were generated using GraphPad Prism and R version 4.6.1, as appropriate.

#### Experimental statistical analysis

Statistical analyses of in vitro and in vivo experiments were performed using GraphPad Prism version 9.3.1. Comparisons between two groups were performed using two-sided unpaired Student’s t tests unless otherwise indicated. Longitudinal measurements and comparisons involving multiple treatment groups were analyzed using two-way ANOVA, as appropriate. Survival curves were estimated using the Kaplan-Meier method and compared using the log-rank test. Data are presented as mean ± standard deviation (SD) or mean ± standard error of the mean (SEM), as indicated in the corresponding figure legends. The numbers of biological replicates or animals are specified in the figure legends. All statistical tests were two-sided, and *P* < 0.05 was considered statistically significant.

