## Supplemental Information for "NECTIN4 defines a subtype-dependent tumor state coupling enfortumab vedotin targetability to antitumor immunity in bladder cancer"

### Figure S1

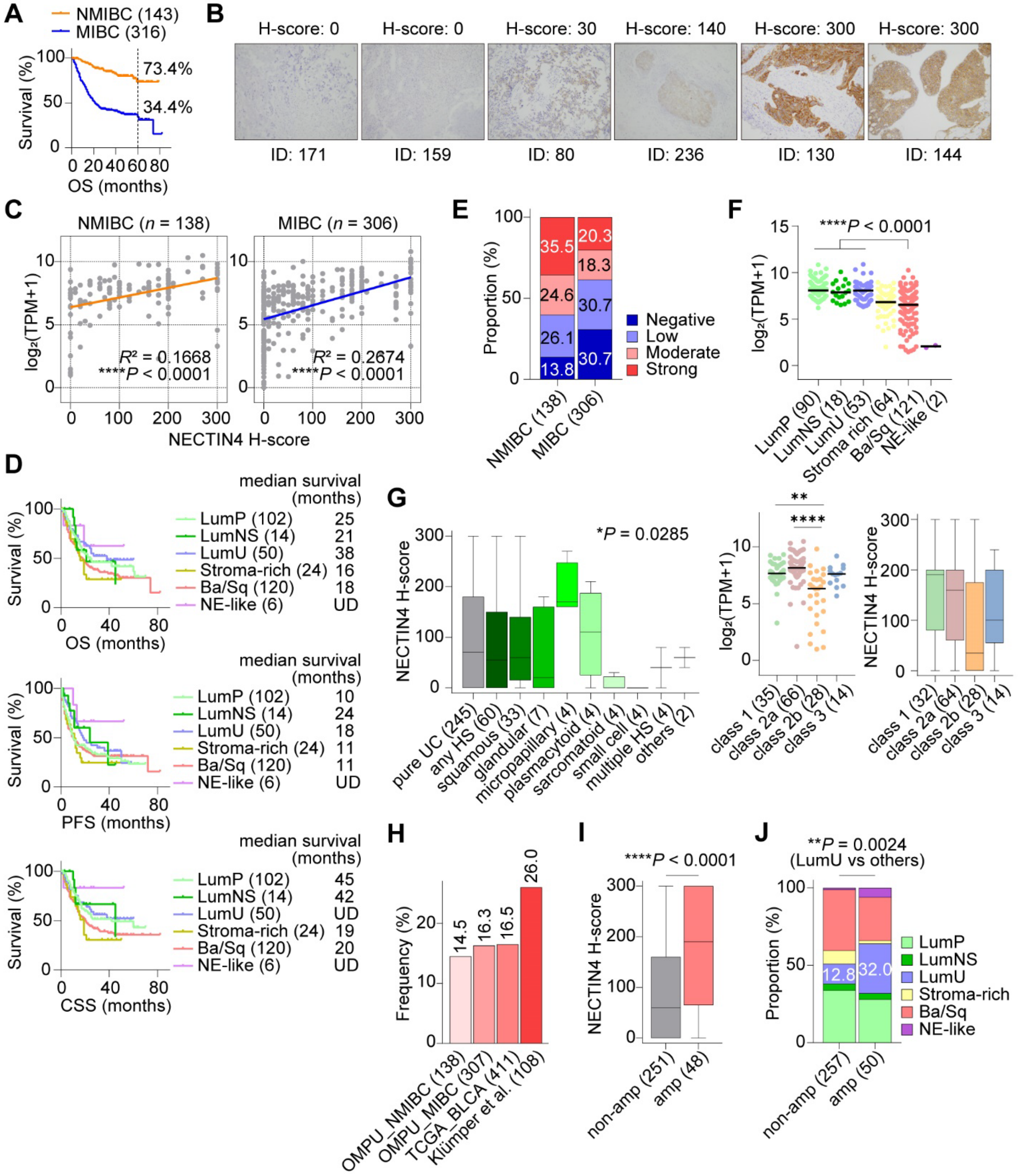


**Figure S1. Clinicopathologic and genomic features associated with NECTIN4 expression in bladder cancer (BC), related to Figure 1**

Unless otherwise indicated, analyses were performed in the OMPU-NCC cohort.
(A) Kaplan-Meier analysis of overall survival (OS) in non-muscle-invasive bladder cancer (NMIBC) and muscle-invasive bladder cancer (MIBC). Five-year OS rates are indicated.
(B) NECTIN4 immunohistochemistry (IHC). Original magnification, ×200.
(C) Correlations between NECTIN4 mRNA expression and H-score in NMIBC and MIBC.
(D) Kaplan-Meier analyses of OS, progression-free survival (PFS), and cancer-specific survival (CSS) across consensus molecular subtypes of MIBC.
(E) Distribution of NECTIN4 IHC categories in NMIBC and MIBC.
(F) NECTIN4 mRNA expression across consensus molecular subtypes in the IMvigor210 cohort (top), and NECTIN4 mRNA expression and IHC H-scores across UROMOL 2021 classes of NMIBC in the OMPU-NCC cohort (bottom).
(G) NECTIN4 H-scores across histologic subtypes (HSs) of BC.
(H) Frequencies of NECTIN4 amplification.
(I) NECTIN4 H-scores in non-amplified and NECTIN4-amplified MIBCs.
(J) Consensus molecular subtype distributions according to NECTIN4 amplification status in MIBC.
Horizontal lines indicate medians. Boxplots show medians, interquartile ranges, and minimum-to-maximum whiskers. Numbers indicate tumors. *P* values were determined by simple linear regression in (C), Kruskal-Wallis tests in (F) and (G), with Dunn's multiple-comparisons test for pairwise comparisons in (F), Mann-Whitney U tests in (I), and Fisher's exact test in (J). *$P < 0.05$, **$P < 0.01$, and ****$P < 0.0001$. TPM, transcripts per million; UC, urothelial carcinoma; UD, undetermined.

#### Figure S2

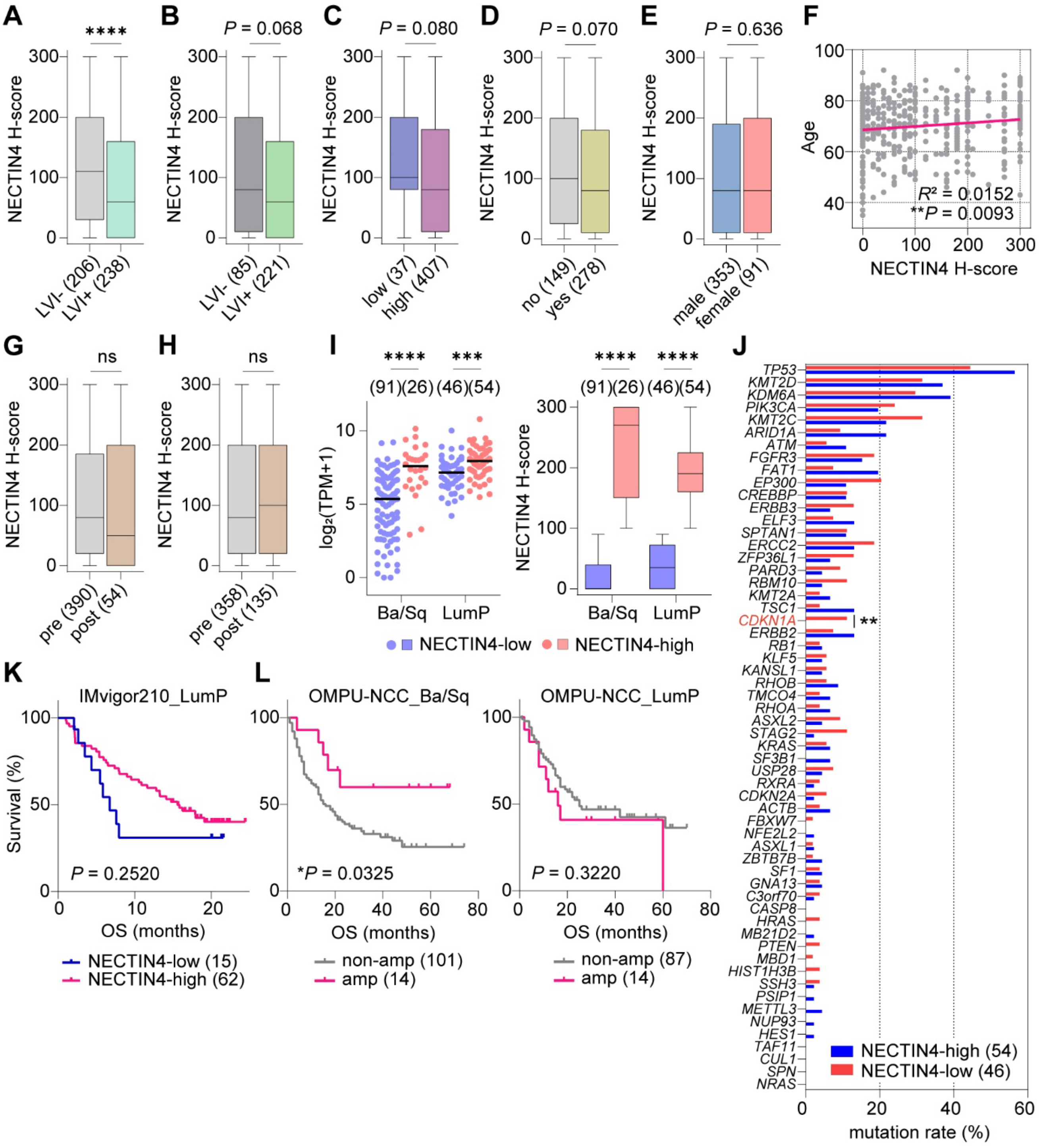

**Figure S2. Clinicopathologic and subtype-specific associations of NECTIN4 expression, related to Figure 2**

Unless otherwise indicated, analyses were performed in the OMPU-NCC cohort.

(A-E) NECTIN4 H-scores according to lymphovascular invasion status in all bladder cancers (A)

and MIBCs (B), tumor grade (C), smoking history (D), and sex (E).
(F) Correlation between age and NECTIN4 H-score.
(G-H) NECTIN4 H-scores before and after radiotherapy (G) or chemotherapy (H).
(I) NECTIN4 mRNA expression and IHC H-scores in Ba/Sq and LumP tumors.
(J) Mutation frequencies in NECTIN4-low and -high LumP MIBCs.
(K) Kaplan-Meier analysis of survival in LumP tumors from the IMvigor210 cohort.
(L) Kaplan-Meier analyses of overall survival (OS) according to NECTIN4 amplification status in Ba/Sq and LumP MIBCs.
Horizontal lines in the scatterplot indicate medians. Boxplots show medians, interquartile ranges, and minimum-to-maximum whiskers. Numbers indicate patients. *P* values were determined by Mann-Whitney U tests in (A-E) and (G-I), simple linear regression in (F), Fisher's exact test in (J), and log-rank tests in (K) and (L). *$P < 0.05$, **$P < 0.01$, ***$P < 0.001$ and ****$P < 0.0001$; ns, not significant. LVI, lymphovascular invasion; amp, amplified; non-amp, non-amplified.

#### Figure S3

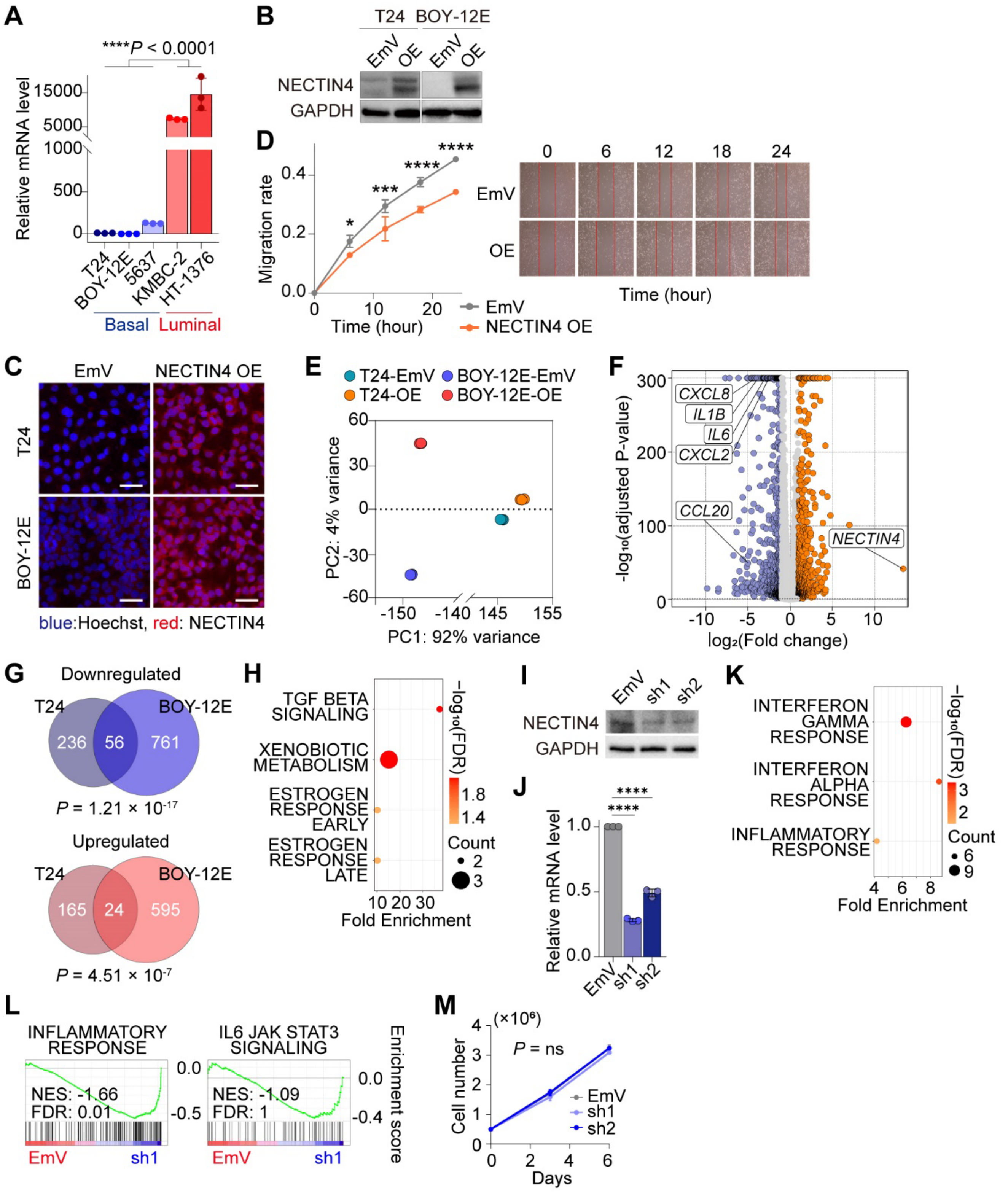


**Figure S3. NECTIN4 expression and subtype-dependent transcriptional effects in bladder**

**cancer (BC) cells, related to Figure 3**

(A) NECTIN4 mRNA expression in basal and luminal BC cell lines.

(B-C) Immunoblot (B) and immunofluorescence (C) analyses of NECTIN4. Scale bar, 100 μm.

(D) Scratch migration assay in BOY-12E cells.

(E) Principal-component analysis of transcriptomic profiles.

(F) Differentially expressed genes between NECTIN4-overexpressing (OE) and empty vector (EmV) BOY-12E cells.

(G) Overlap of genes downregulated or upregulated by NECTIN4 overexpression.

(H) Hallmark pathways enriched among genes commonly upregulated by NECTIN4 overexpression.

(I-J) Immunoblot (I) and RT-qPCR (J) validation of NECTIN4 knockdown (KD) in KMBC-2 cells.

(K) Hallmark pathways enriched among genes upregulated following NECTIN4 KD.

(L) Gene set enrichment analysis of inflammatory response and IL6-JAK-STAT3 signaling following NECTIN4 KD.

(M) Cell proliferation following NECTIN4 KD.

Data in (A), (D), (J), and (M) are mean ± SD; $n = 3$. *P* values were determined by two-sided unpaired t tests in (A) and (J), two-way ANOVA in (D) and (M), and hypergeometric test in (G). *$P < 0.05$, ***$P < 0.001$ and ****$P < 0.0001$; ns, not significant. FDR, false discovery rate; NES, normalized enrichment score.

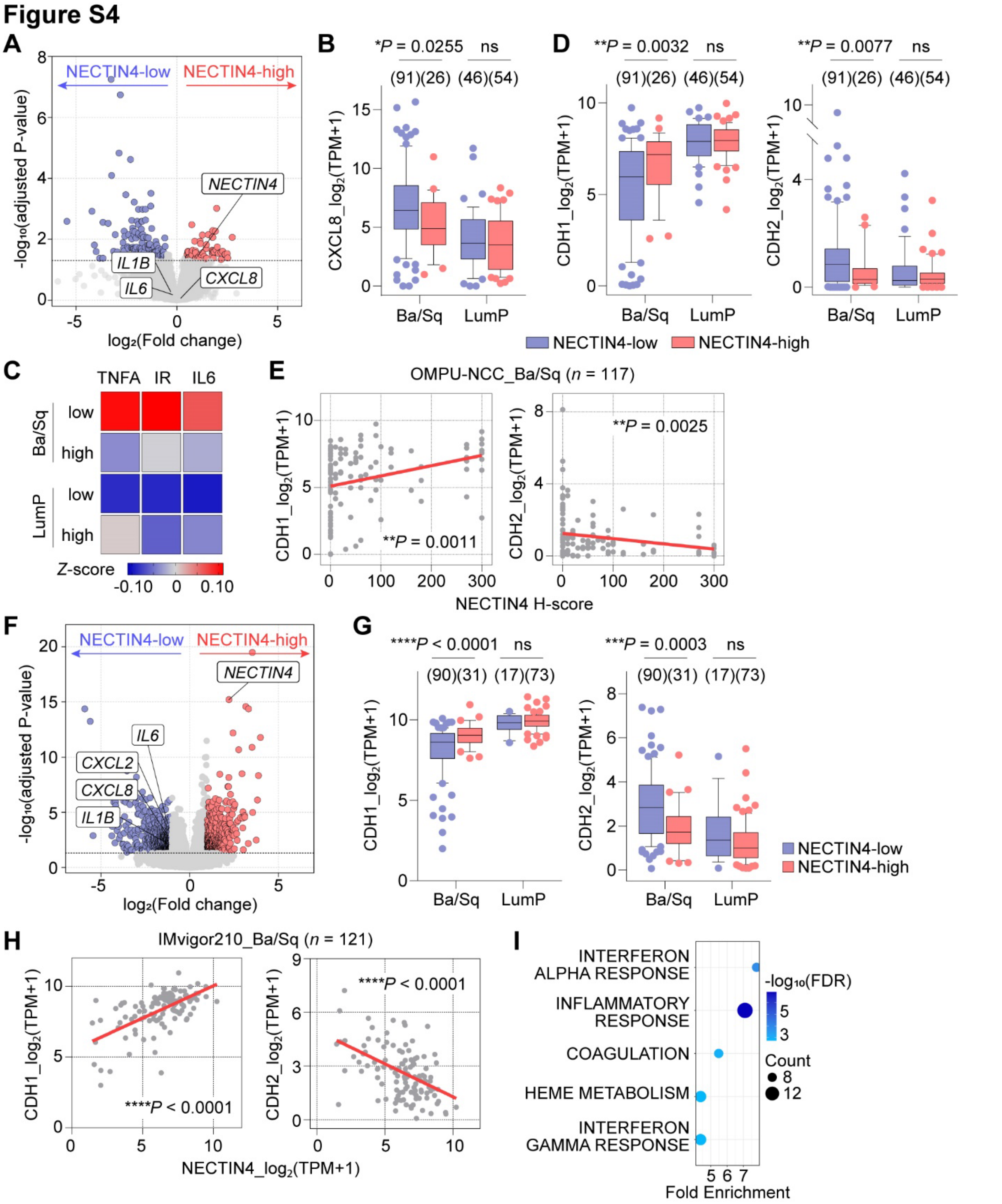


**Figure S4. Subtype-specific inflammatory and epithelial–mesenchymal programs associated with NECTIN4 expression, related to Figure 4**

(A-E) Analyses of the OMPU-NCC cohort.

(A) Differentially expressed genes (DEGs) between NECTIN4-low and -high LumP MIBCs.

(B) CXCL8 expression according to NECTIN4 status in Ba/Sq and LumP tumors.

(C) Mean Hallmark pathway z-scores for TNFα signaling via NF-κB (TNFA), inflammatory response (IR) and IL6-JAK-STAT3 (IL6) signaling according to NECTIN4 status and subtype.

(D) CDH1 and CDH2 expression according to NECTIN4 status and subtype.

(E) Correlations between NECTIN4 H-score and CDH1 or CDH2 expression in Ba/Sq tumors.

(F-I) Analyses of the IMvigor210 cohort.

(F) DEGs between NECTIN4-low and -high Ba/Sq tumors.

(G) CDH1 and CDH2 expression according to NECTIN4 status and subtype.

(H) Correlations between NECTIN4 and CDH1 or CDH2 expression in Ba/Sq tumors.

(I) Top five Hallmark pathways enriched in NECTIN4-low LumP tumors.

Numbers indicate patients. $P$ values were determined by Mann-Whitney U tests in (B), (D), and (G), simple linear regression in (E) and (H). $*P < 0.05$, $**P < 0.01$, $***P < 0.001$ and $****P < 0.0001$; ns, not significant. FDR, false discovery rate.

**Figure S5**

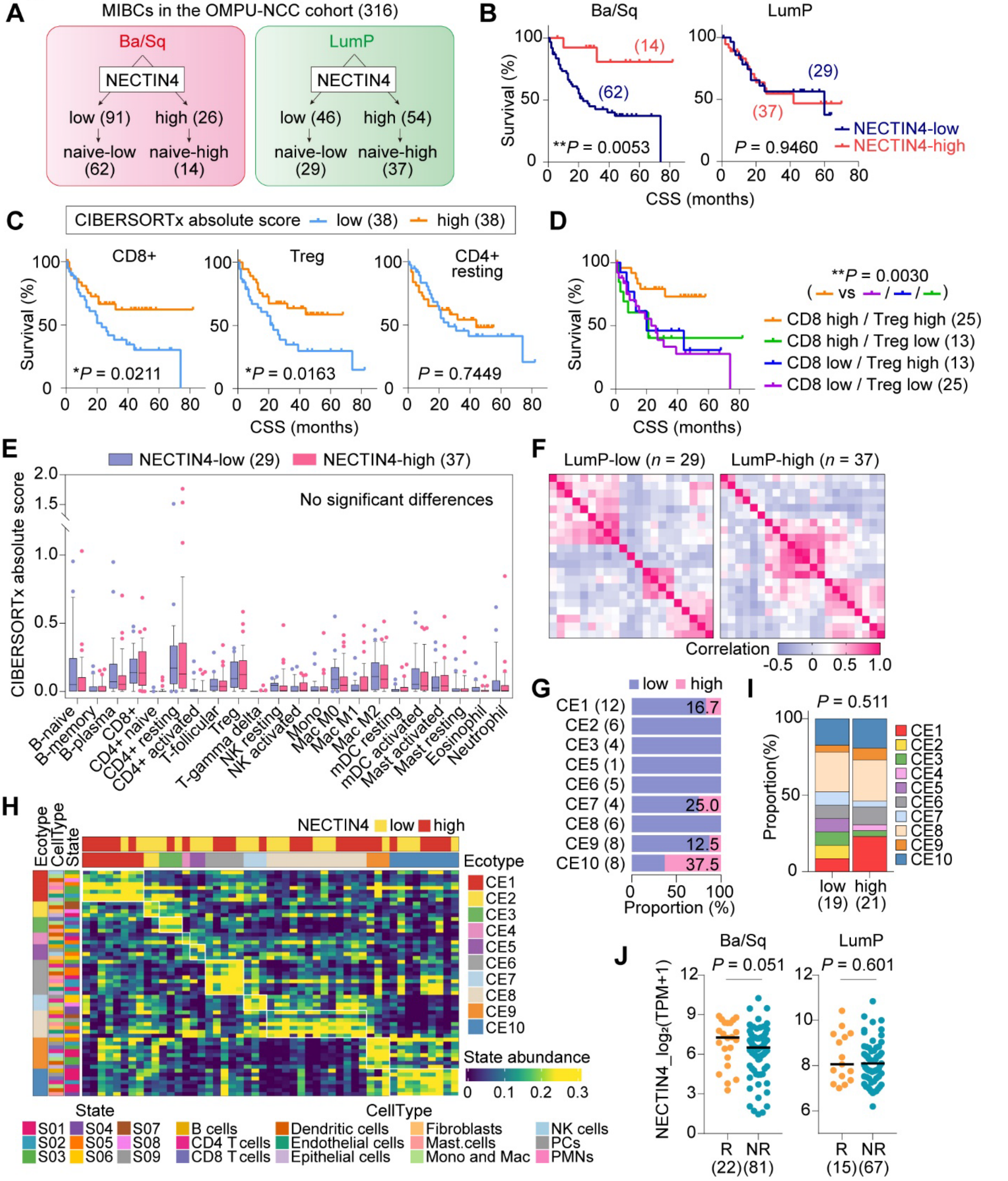


**Figure S5. Subtype-specific immune features associated with NECTIN4 expression and response to immune checkpoint inhibitors (ICIs), related to Figure 5**

Unless otherwise indicated, analyses were performed in treatment-naïve MIBCs from the OMPU-NCC cohort.

(A) Selection of treatment-naïve Ba/Sq and LumP MIBCs according to NECTIN4 expression.

(B) Kaplan-Meier analyses of cancer-specific survival (CSS) in Ba/Sq and LumP tumors.

(C) Kaplan-Meier analyses of CSS according to CIBERSORTx-inferred abundance of CD8+ T cells, regulatory T cells (Tregs), and resting memory CD4+ T cells (CD4+ resting) in Ba/Sq tumors. Patients were dichotomized into low- and high-abundance groups using the median value for each cell type.

(D) Kaplan-Meier analysis of CSS according to combined CD8+ T-cell and Treg abundance in Ba/Sq tumors. CD8-high/low and Treg-high/low groups were defined using CIBERSORTx absolute-score cutoffs of 0.18 and 0.04, respectively.

(E-F) CIBERSORTx-inferred immune-cell abundance (E) and Pearson correlation matrices (F) in NECTIN4-low and -high LumP tumors.

(G) Distribution of NECTIN4-low and -high tumors across cell ecotypes (CEs) in Ba/Sq MIBC.

(H-I) EcoTyper-derived cell states and CE assignments (H) and CE distributions according to NECTIN4 status (I) in LumP tumors.

(J) NECTIN4 expression in ICI responders (R) and nonresponders (NR) with Ba/Sq or LumP tumors in the IMvigor210 cohort.

Responders were defined as patients with complete or partial response and nonresponders as those with stable or progressive disease. Numbers indicate patients. Horizontal lines indicate medians. Boxplots show medians, interquartile ranges, and 10th-90th percentile whiskers. *P* values were determined by log-rank tests in (B-D), two-way ANOVA in (E), Fisher's exact test in (I), and Mann-Whitney U test in (J). **P* < 0.05 and ***P* < 0.01.

#### Figure S6

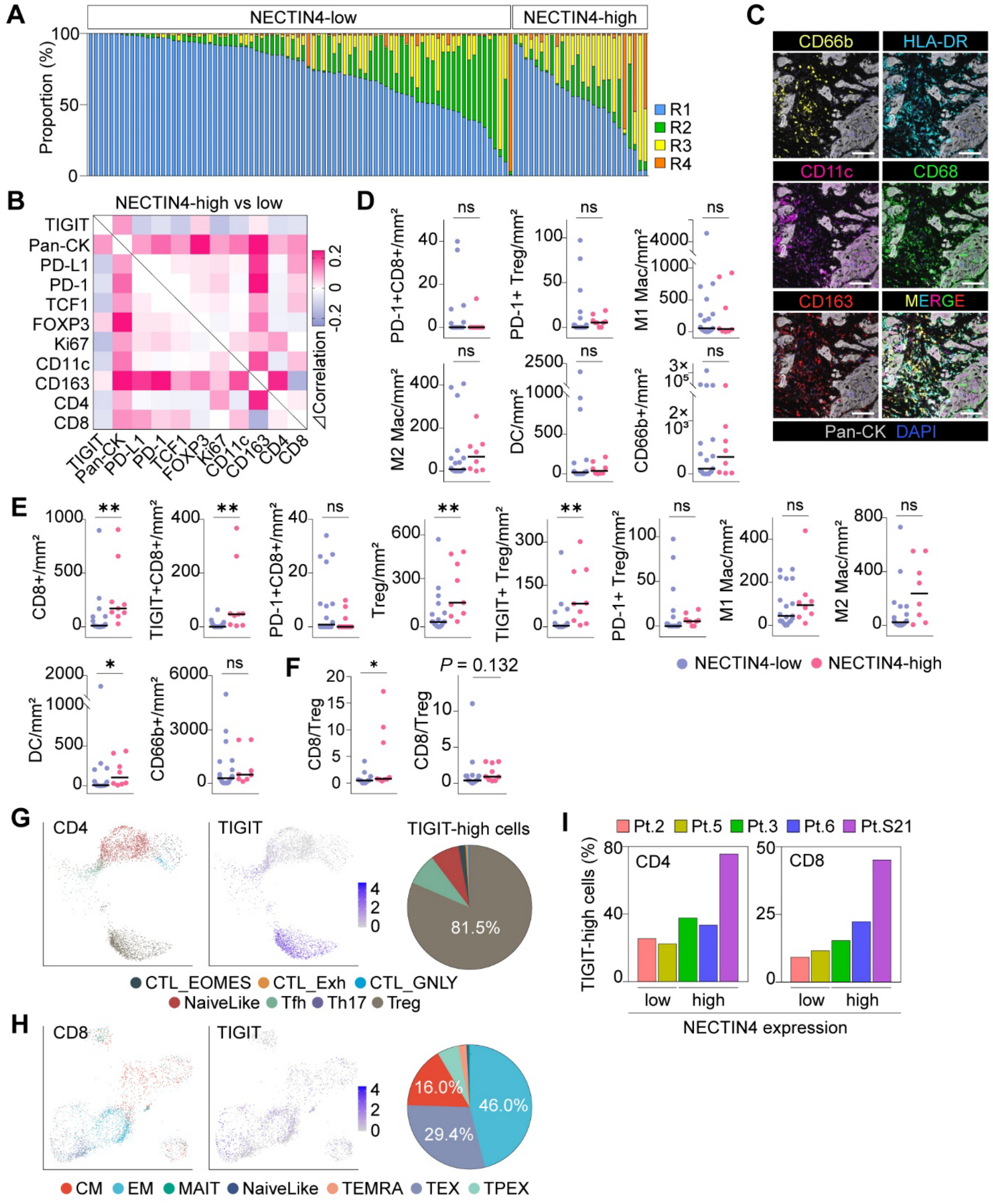


**Figure S6. Spatial and single-cell immune features associated with NECTIN4 expression in Ba/Sq MIBC, related to Figure 6**

(A) Proportions of the four CytoMAP-defined regions across individual CODEX-profiled Ba/Sq tumors.

(B) Differences in pairwise correlations among the indicated tumor and immune features in NECTIN4-high versus -low tumors. Positive values indicate stronger correlations in NECTIN4-high tumors.

(C) Opal-based mIF images from the myeloid-cell panel. Scale bars, 100 μm.

(D-E) Densities of the indicated immune-cell populations in tumor (D) and stromal (E) compartments.

(F) Intratumoral (left) and stromal (right) CD8/Treg ratios in NECTIN4-low and -high tumors.

(G-H) UMAPs of CD4+ (G) and CD8+ (H) T cells mapped onto the reference TIL atlas, colored by cell state or TIGIT expression. Pie charts show the composition of TIGIT-high cells across T-cell states.

(I) Proportions of TIGIT-high cells among CD4+ and CD8+ T cells in NECTIN4-low and -high tumors.

Horizontal lines indicate medians. *P* values were determined by Mann-Whitney U tests in (D-F). **P* < 0.05 and ***P* < 0.01; ns, not significant. DC, dendritic cell; Mac, macrophage; Pan-CK, pan-cytokeratin; Treg, regulatory T cell; FC, fold enrichment; TIL, tumor-infiltrating lymphocyte; CTL, cytotoxic T lymphocyte; Tfh, T follicular helper; Th17, T helper 17; CM, central memory; EM, effector memory; MAIT, mucosal-associated invariant T; TEMRA, terminally differentiated effector memory re-expressing CD45RA; TEX, exhausted T cell; TPEX, progenitor exhausted T cell.

#### Figure S7

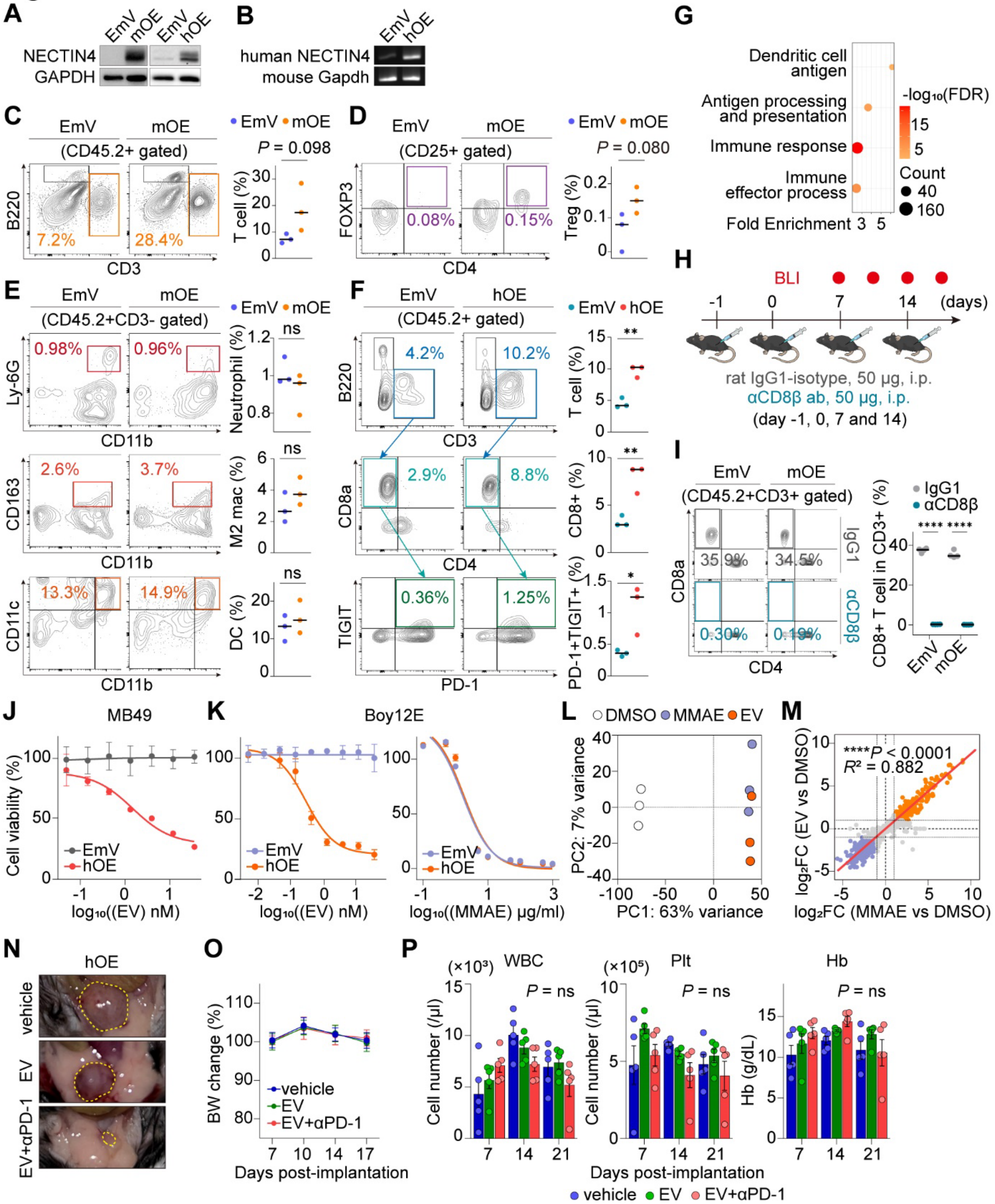


**Figure S7. Immune phenotypes and EV targetability in Nectin4/NECTIN4-expressing orthotopic bladder tumors, related to Figure 7**

(A-B) Immunoblot analysis of murine and human NECTIN4 (A) and RT-PCR analysis of human NECTIN4 (B) in empty-vector (EmV), murine Nectin4-overexpressing (mOE), or human NECTIN4-overexpressing (hOE) MB49 cells.

(C-E) Flow-cytometric quantification of CD3+ T cells (C), regulatory T cells (Tregs) (D), and neutrophils, M2-like macrophages, and dendritic cells (DCs) (E) in orthotopic EmV and mOE tumors. $n = 3$.

(F) Flow-cytometric quantification of total T cells, CD8+ T cells, and PD-1+TIGIT+CD8+ T cells in orthotopic EmV and hOE tumors. $n = 3$.

(G) Selected immune-related pathways enriched in CD45.2+ immune cells isolated from hOE versus EmV tumors.

(H-I) Experimental schema (H) and flow-cytometric confirmation (I) of systemic CD8+ T-cell depletion in mice bearing orthotopic EmV or mOE tumors. $n = 3$ in (I).

(J) Viability of EmV and hOE MB49 cells treated with enfortumab vedotin (EV).

(K) Viability of EmV and hOE BOY-12E cells treated with EV or monomethyl auristatin E (MMAE).

(L-M) Principal-component analysis (L) and correlation of transcriptomic responses to MMAE and EV relative to DMSO (M) in hOE BOY-12E cells.

(N-P) Bladder images (N), body weight (BW) (O), and peripheral blood counts (P) in mice bearing hOE tumors treated with vehicle, EV, or EV+anti-PD-1. $n = 5$ in (O) and (P).

Data in (O) and (P) are mean ± SD. Horizontal lines indicate medians. $P$ values were determined by two-sided unpaired t tests in (C-F) and (I) and at each time point in (P). $^{*}P < 0.05$, $^{**}P < 0.01$, $^{***}P < 0.001$ and $^{****}P < 0.0001$; ns, not significant.

#### Figure S8

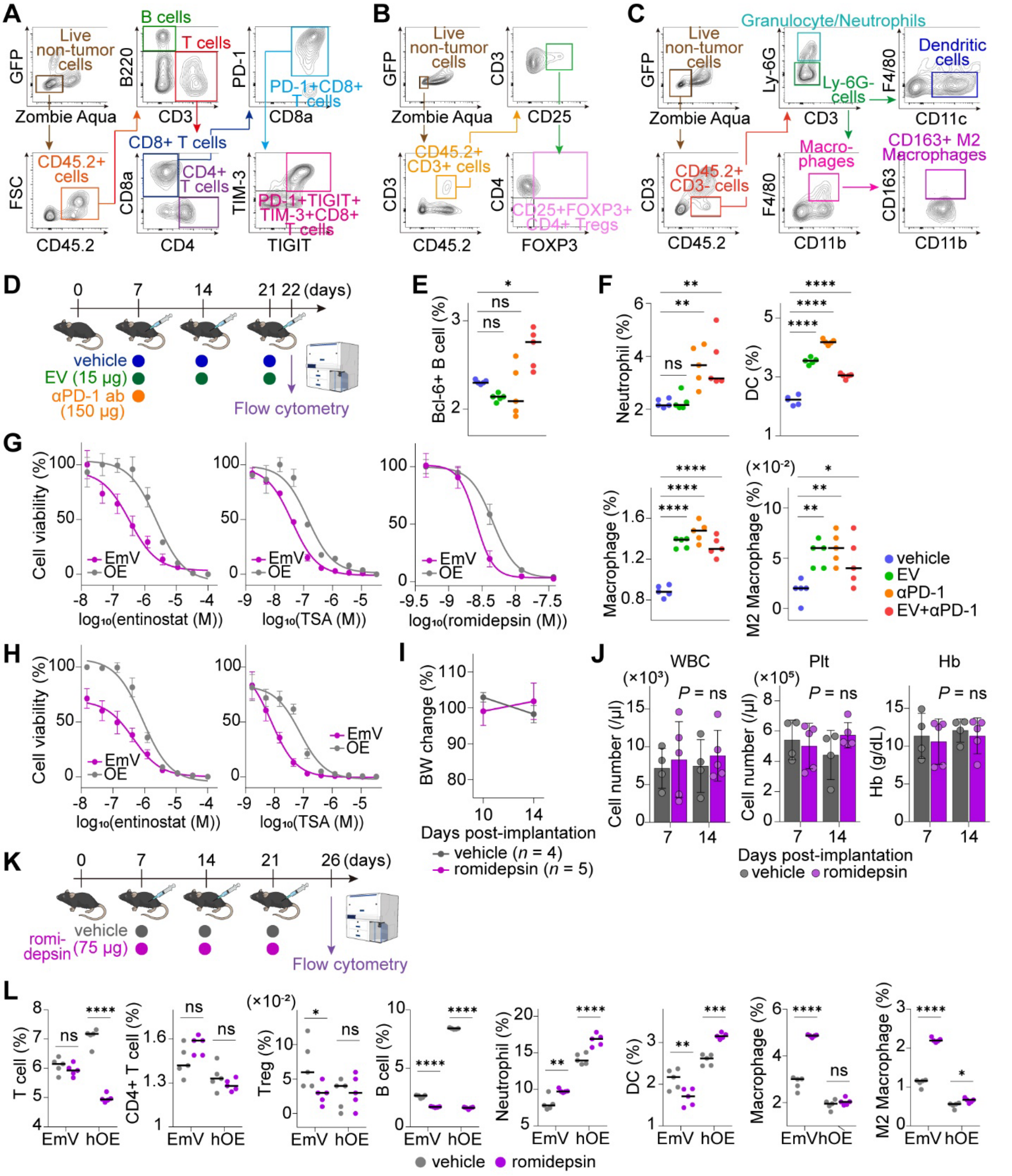


**Figure S8. TME remodeling by EV plus PD-1 blockade and HDAC inhibition, related to Figure 8**

(A-C) Flow-cytometry gating strategies for lymphoid-cell (A), Treg (B), and myeloid-cell (C) profiling of orthotopic tumors.

(D) Experimental schema for flow-cytometric profiling of orthotopic hOE tumors treated with vehicle, enfortumab vedotin (EV), anti-PD-1 antibody (αPD-1), or EV+αPD-1.

(E-F) Frequencies of BCL6+ B cells (E) and the indicated myeloid-cell populations (F) following treatment. $n = 5$.

(G-H) Dose-response curves for entinostat, trichostatin A (TSA) and romidepsin in empty-vector (EmV) and NECTIN4-overexpressing (OE) T24 (G) and BOY-12E (H) cells.

(I-J) Body weight (BW) changes of mice bearing EmV tumors (I) and peripheral white blood cell (WBC) counts, platelet counts (Plt) and hemoglobin (Hb) concentrations (J) during vehicle or romidepsin treatment. Mean ± SD, $n = 4$ or 5.

(K) Experimental schema for flow-cytometric profiling of orthotopic EmV and hOE tumors treated with vehicle or romidepsin.

(L) Frequencies of the indicated lymphoid and myeloid populations following vehicle or romidepsin treatment. $n = 5$.

Horizontal lines indicate medians. $P$ values were determined by one-way ANOVA followed by Tukey's multiple-comparisons test in (E) and (F), two-sided unpaired t tests in (J), and two-way ANOVA followed by Šídák's multiple-comparisons test in (L). *$P < 0.05$, **$P < 0.01$, ***$P < 0.001$ and ****$P < 0.0001$; ns, not significant. DC, dendritic cell; Treg, regulatory T cell.

**Table S1. Clinicopathologic characteristics of patients with Ba/Sq or LumP MIBC stratified by NECTIN4 H-score in the OMPU-NCC cohort**

| | BaSq subtype (*n* = 117) | | *P* | LumP subtype (*n* = 100) | | *P* |
|---|---|---|---|---|---|---|
| NECTIN4 H-score | <100 (*n* = 91) | ≥100 (*n* = 26) | | <100 (*n* = 46) | ≥100 (*n* = 54) | |
| Sex (%) | | | | | | |
| Male | 64 (70.3) | 22 (84.6) | | 38 (82.6) | 42 (77.8) | |
| Female | 27 (29.7) | 4 (15.4) | 0.146 | 8 (17.4) | 12 (22.2) | 0.547 |
| Age | | | | | | |
| median [IQR] | 67[58-77] | 70[59.5- | 0.553 | 72.5 [62.75- | 70 [63- | 0.656 |
| Smoking history (%) | | | | | | |
| never | 32 (35.2) | 9 (34.6) | | 12 (26.1) | 21 (38.9) | |
| former/current | 52 (57.1) | 16 (61.5) | | 34 (73.9) | 31 (57.4) | |
| unknown | 7 (7.7) | 1 (3.8) | 0.776 | 0 (0) | 2 (3.7) | 0.137 |
| TNM classification (%) | | | | | | |
| cN0M0 | 64 (70.3) | 19 (73.1) | | 33 (71.7) | 37 (68.5) | |
| cN1M0 | 19 (20.9) | 3 (11.5) | | 7 (15.2) | 10 (18.5) | |
| cNxM1 | 8 (8.8) | 4 (15.4) | 0.403 | 6 (13.0) | 7 (13.0) | 0.907 |
| Concomitant CIS (%) | | | | | | |
| no | 85 (93.4) | 24 (92.3) | | 43 (93.5) | 47 (87.0) | |
| yes | 6 (6.6) | 2 (7.7) | 0.845 | 3 (6.5) | 7 (13.0) | 0.285 |
| Lymphovascular invasion | | | | | | |
| no | 22 (24.2) | 5 (19.2) | | 16 (34.8) | 19 (35.2) | |
| yes | 69 (75.8) | 21 (80.8) | 0.598 | 30 (65.2) | 35 (64.8) | 0.966 |
| Histological variants (%) | | | | | | |
| no | 64 (70.3) | 19 (73.1) | | 42 (91.3) | 45 (83.3) | |
| yes | 27 (29.7) | 7 (26.9) | 0.786 | 4 (8.7) | 9 (16.7) | 0.238 |
| Previous BCG treatment (%) | | | | | | |
| no | 85 (93.4) | 20 (76.9) | | 41 (89.1) | 50 (92.6) | |
| yes | 6 (6.6) | 6 (23.1) | 0.015* | 5 (10.9) | 4 (7.4) | 0.547 |
| Previous chemotherapy (%) | | | | | | |
| no | 66 (72.5) | 17 (65.4) | | 34 (73.9) | 39 (72.2) | |
| yes | 25 (27.5) | 9 (34.6) | 0.44 | 12 (26.1) | 15 (27.8) | 0.849 |
| Previous radiation therapy | | | | | | |
| no | 73 (80.2) | 20 (76.9) | | 39 (84.8) | 44 (81.5) | |
| yes | 18 (19.8) | 6 (23.1) | 0.714 | 7 (15.2) | 10 (18.5) | 0.661 |
| Follow-up period | | | | | | |
| median [IQR] (months) | 13 [5-26] | 22 [6-42.5] | 0.047* | 17 [9-46.25] | 17 [9.75-27] | 0.31 |

Ba/Sq, basal/squamous; BCG, bacillus Calmette–Guérin; CIS, carcinoma in situ; IQR, interquartile range; LumP, luminal papillary; MIBC, muscle-invasive bladder cancer; OMPU-NCC, Osaka Medical and Pharmaceutical University–National Cancer Center Japan Consortium; TNM, tumor-node-metastasis. *$P < 0.05$.

**Table S2. Pearson correlation matrices of CIBERSORTx-inferred immune-cell abundance in treatment naïve MIBC tumors**

Data are provided in the accompanying Excel file.

#### Table S3. CODEX/Phenocycler phenotype definitions

| Category | Label | Definitions | Phenotype | Panels |
|---|---|---|---|---|
| Tumor | NECTIN4+ tumor | NECTIN4: Pan-Cytokeratin | NECTIN4-positive epithelial tumor cells | Fig. 6B and 6E |
| Tumor | NECTIN4− tumor | Pan-Cytokeratin (NECTIN4 negative) | NECTIN4-negative epithelial tumor cells | Fig. 6B |
| Stromal | Vascular | CD31 | Endothelial / vascular cells | Fig. 6B |
| Stromal | Podoplanin | Podoplanin | Podoplanin-positive stromal cells | Fig. 6B |
| Stromal | αSMA/ACTA2 | αSMA/ACTA2 | Myofibroblast / CAF-like stromal cells | Fig. 6B |
| Marker-defined | PD-L1+ | PD-L1 | PD-L1-expressing cells | Fig. 6B |
| T cell | PD-1+TCF1+CD8+ | PD-1: TCF1: CD8 | Stem-like / progenitor-exhausted CD8 T cells | Fig. 6B and 6E |
| T cell | TIGIT+CD8+ | TIGIT: CD8 | TIGIT-expressing CD8 T cells | Fig. 6B |
| T cell | TCF1+CD8+ | TCF1: CD8 | TCF1-positive stem-like / memory CD8 T cells | Fig. 6B |
| T cell | PD-1+TIM3+CD8+ | PD-1: TIM3: CD8 | Terminally exhausted CD8 T cells | Fig. 6B |
| T cell | PD-1+TIGIT+CD8+ | PD-1: TIGIT: CD8 | Dysfunctional / exhausted CD8 T cells | Fig. 6B and 6E |
| T cell | PD-1+CD8+ | PD-1: CD8 | PD-1-expressing CD8 T cells | Fig. 6B and 6E |
| T cell | CD8+ | CD8 | CD8 T cells | Fig. 6B and 6E |
| T cell | CXCL13+CD4+ | CXCL13: CD4 | CXCL13-producing CD4 T cells | Fig. 6B and 6E |
| T cell | Treg | FOXP3: CD25: CD4 | Regulatory T cells | Fig. 6B and 6E |
| T cell | CD4+ | CD4 | CD4+ T cells | Fig. 6B |
| B / plasma | CD38+ plasma | CD38 | Plasma cells | Fig. 6B |
| B cell | CD20+CD21− | CD20 (CD21 negative) | CD21-negative B cells | Fig. 6B |
| Myeloid | CD163+ | CD163 | CD163-positive macrophage / myeloid cells | Fig. 6B |
| Myeloid | CD68+ | CD68 | Macrophages | Fig. 6B |
| Myeloid | CD68+CD163+ | CD68: CD163 | M2-like macrophages | Fig. 6B |
| Myeloid | HLA-DR+CD14+ | CD14:HLA-DR | HLA-DR-positive monocytes / myeloid cells | Fig. 6B and 6E |
| Myeloid / DC | HLA-DR+CD11c+ | CD11c:HLA-DR | HLA-DR-positive dendritic cells | Fig. 6B and 6E |
